# Absence of a prolonged macrophage and B cell response inhibits heart regeneration in *the Mexican cavefish*

**DOI:** 10.1101/2025.04.24.650152

**Authors:** Esra Sengul, Helen G. Potts, William T. Stockdale, Ryan D. Carter, Laura Bevan, Maria Nozdrina, Rita Alonaizan, Zhilian Hu, Abigail Goodship, Jun Ying, Konstantinos Lekkos, Lucy O’Byrne, Madeleine E. Lemieux, Rebecca Richardson, Mathilda T.M. Mommersteeg

**Affiliations:** Institute of Developmental and Regenerative Medicine, Old Road Campus, University of Oxford, Oxford, UK; Department of Physiology, Anatomy & Genetics, University of Oxford, Oxford, UK; School of Physiology, Pharmacology and Neuroscience, Faculty of Health and Life Sciences, University of Bristol, University Walk, Bristol BS8 1TD, UK; Bioinfo, Plantagenet, ON, Canada

## Abstract

A balanced immune response after cardiac injury is crucial to successful heart regeneration, but knowledge of what distinguishes a regenerative from a scarring response is still limited. The Mexican cavefish provides a unique comparative model to study heart regeneration and scarring within a single species. Surface-dwelling fish are capable of heart regeneration whereas their cave-dwelling Pachón counterparts lack this ability, similar to the human heart. Using single-cell transcriptomics and immune perturbations, we find significant differences in the immune response between the two populations. Unlike the transient response in the scarring Pachón, the regenerative surface fish heart generates an unexpected functionally active prolonged innate and adaptive immune response at the late stages of regeneration. Inhibiting the overall prolonged immune response impairs regeneration and cardiomyocyte proliferation. Further characterisation of specific cell types shows that late-present macrophages are phagocytic, and their depletion disrupts regeneration but not cardiomyocyte proliferation while inhibiting B cells impairs regeneration by reducing cardiomyocyte proliferation. This B cell response is conserved in zebrafish. Our findings reveal critical immune mechanisms distinguishing regenerative and non-regenerative responses, offering insights for potential therapeutic strategies to enhance heart repair.

## Introduction

Regeneration of the adult heart is a remarkable ability seen in some species, such as zebrafish, newts, and axolotls^1–4^. This response is absent in humans, where injury to the heart often results in scarring and permanent damage. After myocardial infarction (MI), the human heart undergoes an acute inflammatory response as dying cells release damage-associated molecular patterns (DAMPs), triggering immune cells to release cytokines and chemokines^5^. This cascade activates a host of immune responses aimed at repairing the damaged tissue, but also causing excessive inflammation, extracellular matrix (ECM) remodelling and scar formation rather than regeneration^6,7^. Over time, this fibrotic response can compromise cardiac function and lead to heart failure. However, the immune response has also been shown to be essential for successful regeneration in species such as zebrafish and the neonatal mouse^8,9^.

In both regenerative and non-regenerative species, innate immune cells, such as neutrophils and macrophages, infiltrate the injury site, initiating an acute inflammatory response. However, in regenerative species, these immune components play critical roles in clearing debris, promoting angiogenesis, and stimulating cardiomyocyte proliferation, one of the hallmarks of successful regeneration^10–12^. It is striking how the same immune components can either hinder or support successful regeneration, relying on a delicate, tightly regulated balance. For instance, prolonged neutrophil activity can impair healing, while macrophages can promote either regeneration or drive scarring, depending on their activation state and timing9,13,14.

Adaptive immune cells have also been implicated in heart regeneration, with emerging evidence suggesting that B cells may influence cardiomyocyte proliferation and modulate inflammation, making them an intriguing focus for regenerative studies^15–18^. However, their role in the regenerative process has not yet been fully elucidated, and further research is needed to determine whether targeting B cell activity could enhance cardiac repair. Understanding the immune response that enables successful regeneration in certain species versus scarring in others may provide valuable insights for designing therapeutics to enhance post-MI cardiac repair in humans.

To explore the interplay between immunity and heart regeneration, the Mexican cavefish, *Astyanax mexicanus*, offers a compelling model. This species exists in two main morphs: surface-dwelling fish, capable of regenerating their hearts after injury, and cave-dwelling fish, which have lost this ability and instead form a permanent scar ^19^. The cavefish have evolved in the isolated, nutrient-deficient, and low-diversity environments of Mexico’s underground caves. They adapted to long absence of light and are blind, with altered immune activity due to reduced pathogen exposure^20^. This unique evolutionary background provides a natural framework for studying the genetic and immune differences that influence natural regenerative outcomes without the complications of cross-species comparisons^21^.

In this study, we use a multidimensional approach to dissect the role of immune mechanisms during cardiac regeneration, comparing the surface fish against their cave-dwelling Pachón counterparts. Our findings reveal that surface fish exhibit a prolonged, balanced immune response, characterised by a sustained presence of both myeloid and lymphoid populations, that supports regeneration, while Pachόn fish show a rapid but transient response that culminates in scarring. Inhibition of this prolonged inflammation in surface fish with a broad-acting anti-inflammatory drug, Dexamethasone, results in reduced cardiomyocyte proliferation and impaired regeneration. A detailed characterisation of the immune cell populations between surface fish and Pachón fish shows that the difference in leukocyte numbers is driven by higher numbers of macrophages and B cells in the surface fish during the later recovery stages. Specific depletion of late-stage phagocytic macrophages results in defective scar removal without affecting cardiomyocyte proliferation. B cells in surface fish increase in number from 7-days post cryoinjury (dpci) and pharmacological inhibition of B cells significantly impairs regeneration by reducing cardiomyocyte proliferation, mirroring the defects observed with Dexamethasone treatment.

Collectively, these findings enhance our understanding of the cellular and molecular interplay that underpins cardiac repair and regeneration, pointing to responses of specific immune cells and responses that promote regeneration rather than scarring and could serve as targets for therapeutic interventions in regenerative medicine.

### Inhibition of prolonged inflammation impairs heart regeneration

To characterise the immune response in the regenerative and scarring settings, we performed cryoinjury on surface fish and Pachόn hearts. Ventricles were collected at 1-, 3-, 7-, and 14 dpci, as well as uninjured and 3 days-post sham (dps) controls. Samples were processed for 10X Chromium single-cell RNA-sequencing (scRNA-seq), yielding a sequencing depth of ∼50,000 reads per cell (Fig. 1a). We identified all expected cardiac cell types (Fig. 1b, Supplementary Fig. 1a-b) and examined leukocyte dynamics by sub-setting cells expressing canonical markers *ptprc*, *lcp1* and *cxcr4b* (Supplementary Fig.1c). Plotting the proportion of immune cells relative to other major cardiac cell types revealed that the immune response diverged between regenerative surface fish and scarring Pachόn hearts, particularly at late stages after injury (Fig. 1c). Differential proportion analysis (DPA) showed that Pachόn leukocytes were elevated at 1- and 3-dpci, while surface fish leukocytes were elevated at all time points compared to uninjured levels, including at 7- and 14-dpci (Fig. 1d). We confirmed these scRNA-seq findings on sections using an RNAscope *in situ* hybridisation probe targeting the pan-leukocyte marker *ptprc* (Fig. 1e–f). After an initially similar response, there was a decline in numbers towards baseline in Pachón wounds after 3-dpci. This is expected after the initial peak as observed in other non-regenerative species^6,22^. However, leukocyte numbers remained high in surface fish, with significantly higher numbers than in Pachón at 14-dpci (Fig. 1f), suggesting that prolonged inflammation is a key factor contributing to cardiac repair.

**Fig. 1.**
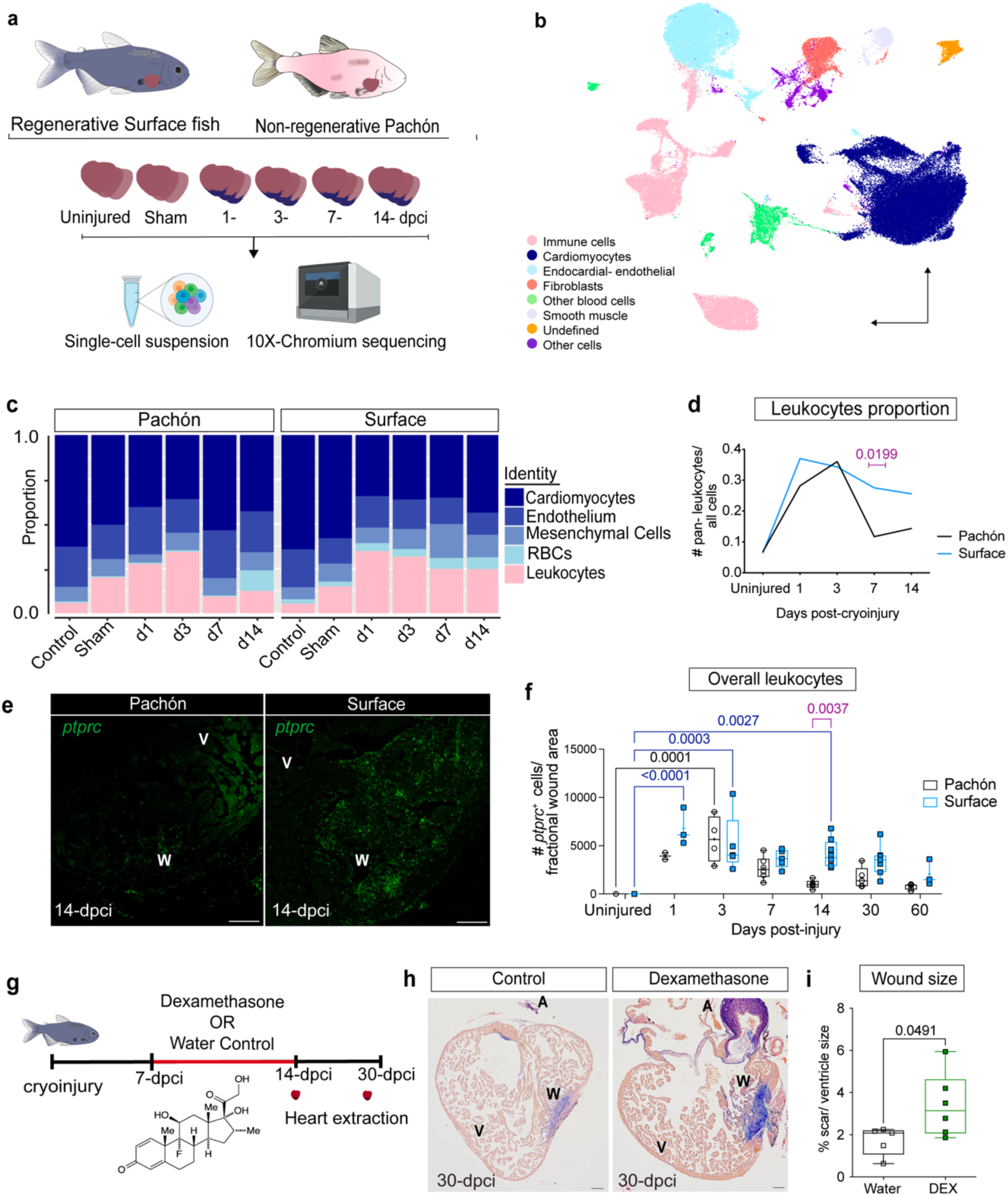
Prolonged inflammation after cardiac injury is required for successful repair in surface fish hearts. **a** Pachόn and surface fish ventricles were collected uninjured, 3 days-post sham (dps), 1-, 3-, 7-, and 14-days post cryoinjury (dpci), dissociated into single cell suspensions and 10X-Chromium single-cell RNA sequencing was performed with a depth of ∼50,000 reads/cell. **b** Annotated UMAP showing all the expected cell types of the heart in the SCT-integration consisting of 85,516 cells. **c** Stacked histogram of major cardiac cell proportions for Pachόn (left) and surface fish (right) hearts at all time points showing that the proportion of Pachόn leukocytes decreases at late time points compared to surface fish. **d** Differential proportion analysis on leukocytes at different time points in Pachón shows higher leukocyte cell numbers at 1- and 3-dpci compared to uninjured with surface fish showing elevated leukocyte numbers at all time points compared to uninjured. **e** RNAscope to reveal *ptprc*^+^ pan-leukocytes in Pachón and surface fish ventricles at 14-dpci, scale bar 100-μm. Insert shows absence of staining for myocardial marker MF20 in the wound region with DAPI labelling for all nuclei. **f** Quantification of *ptprc*^+^ cells present in the wound at 1-, 3-, 7-, 14-, 30-, 60-dpci and uninjured in Pachón and surface fish shows that overall immune response in surface fish and Pachόn hearts significantly diverges at 14-dpci over 60-days’ time course after injury, cell counts were normalised to the fractional wound area, Two-way ANOVA (p adjusted < 0.05 shown, n=3-8 for each fish at all time points). **g** surface fish were exposed to Dexamethasone as water immersion from 7-to 14-dpci after cryoinjury, hearts were collected at 14-dpci and 30-dpci. **h** AFOG-stained images of water control and Dexamethasone-treated surface fish hearts at 30-dpci, scale bar 100-μm. **i** Quantification of the difference in wound area between the control and Dexamethasone-treated ventricles at 30-dpci shows inhibition of prolonged inflammation impairs regeneration in surface fish, p=0.0491 unpaired t-test, n=6 each group. DEX, dexamethasone; dpci, days post cryo-injury; V, ventricle; A, atrium; W, wound.

To test whether this late immune response benefits regeneration, we used the broad-spectrum anti-inflammatory agent Dexamethasone^23^ to suppress the late-stage immune response in surface fish. Fish were treated with Dexamethasone from 7- to 14-dpci (Fig. 1g). Hearts were isolated at 30-dpci, and wound size was analysed using AFOG staining, revealing significantly larger scars in Dexamethasone-treated hearts compared to untreated controls (Fig. 1h–i). These findings highlight the importance of a sustained immune response in promoting effective cardiac repair in surface fish.

### Prolonged innate and adaptive immune responses in surface fish hearts

Since the overall leukocyte response was prolonged in surface fish hearts compared to Pachόn, we further sub-clustered the leukocytes to investigate which immune cell populations were driving this difference (Fig. 2a). We identified the expected leukocyte sub-types with neutrophils, macrophages, and B cells constituting the majority of cells. These populations were present in both surface fish and Pachόn across all time points but decreased at later time points in Pachόn (Fig. 2b). DPA analysis on the different leukocyte sub-types highlighted neutrophils, macrophages and B cells as the most distinct between surface fish and Pachόn (Fig. 2c–e, Supplementary Fig. 2a–e).

**Fig. 2.**
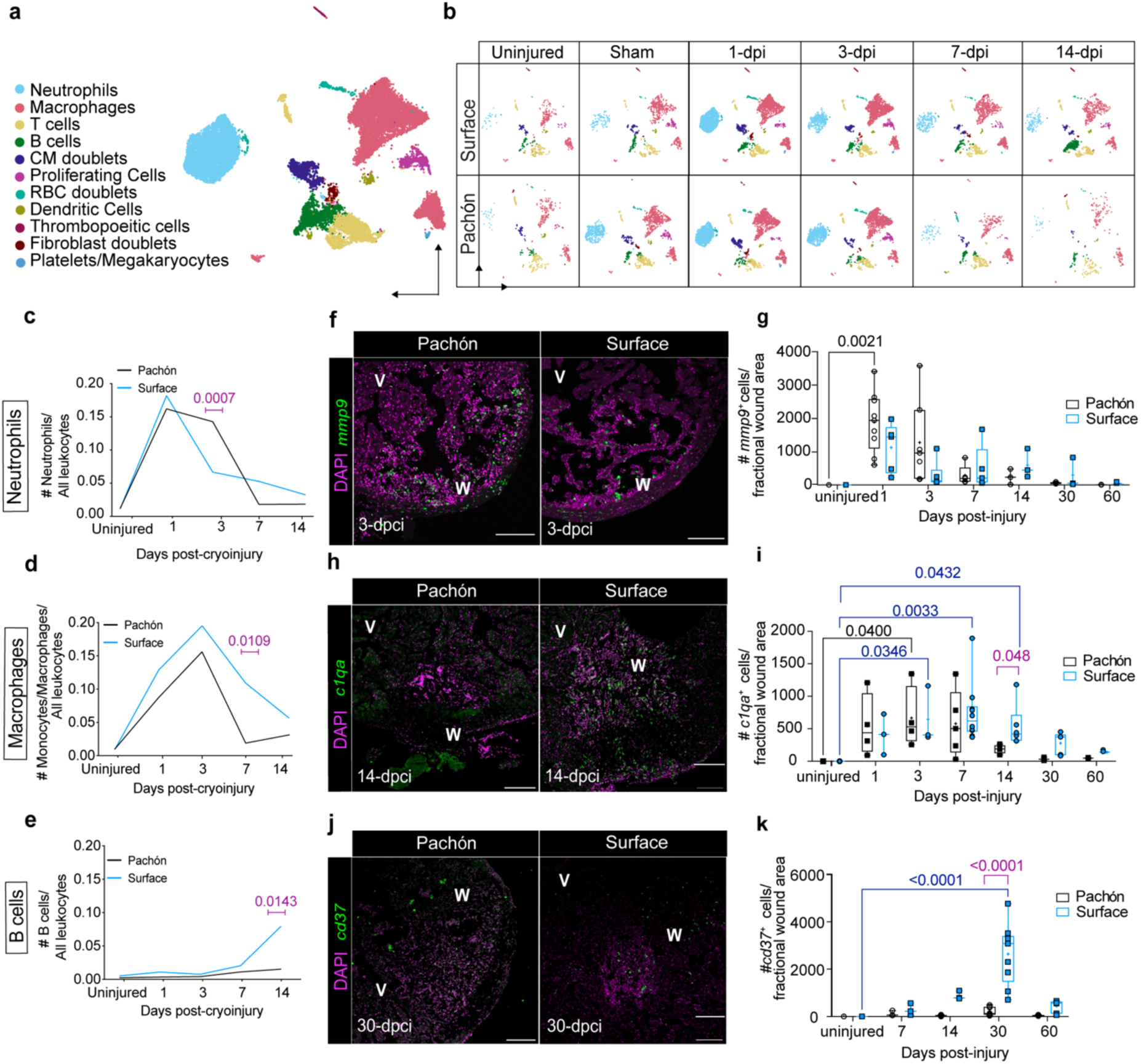
Surface fish has a prolonged innate and adaptive response. **a** Annotated UMAP showing all identified leukocyte types in Pachόn and surface fish heart. **b** UMAPs showing all leukocyte clusters over time in both fish, highlighting that the Pachόn leukocytes captured are decreasing in the late stages of wound healing, but all clusters are still present. **c-e** Differential proportion analysis on neutrophils, monocytes-macrophages and B cells, respectively, in uninjured and at 1-, 3-, 7-, and 14-dpci shows differences between Pachόn and surface fish ventricles. **f** Representative RNAscope images to show *mmp9*^+^ neutrophils at 3-dpci and **g** quantification of *mmp9*^+^ cells in uninjured and at 1-, 3-, 7-, 14-, 30-, 60-dpci in surface fish and Pachόn. **h** Representative RNAscope images to show *c1qa*^+^ macrophages counterstained with DAPI at 14-dpci and **i** quantification of *c1qa*^+^ cells in uninjured and at 1-, 3-, 7-, 14-, 30-, 60-dpci in surface fish and Pachόn. **j** Representative RNAscope images to show *cd37*^+^ B cells at 30-dpci and **k** quantification of *cd37*^+^ cells in uninjured and at 7-, 14-, 30-, 60-dpci in surface fish and Pachόn. Two-way ANOVA, p< 0.05 shown on the graphs, n=3-8 (**g, i, k**). dpci: days post cryo-injury.

We then focused on immune populations that showed the greatest differences in the DPA. Neutrophils, among the first immune cells to arrive at the site of injury, peaked at 3-dpci in both fish populations. However, neutrophil numbers were reduced in Pachόn at later stages, while they remained elevated in surface fish (Fig. 2c). The timely resolution of neutrophils is vital for proper wound healing and for preventing chronic inflammation^22,24,25^, which can lead to impaired healing or fibrosis as we observed in Pachόn. RNAscope for neutrophil marker *mmp9* confirmed the strong initial influx of neutrophils into the wound at 1-dpci. However, in contrast to the single-cell data, there were no significant differences between the fish (Fig. 2f– g).

Macrophages peaked similarly at 3-dpci in both Pachόn and surface fish (Fig. 2d). However, surface fish showed higher macrophage levels compared to Pachόn at 7-dpci, suggesting that macrophages are retained at late stages in regenerating hearts. RNAscope analysis for macrophage marker *c1qa* confirmed that surface fish exhibited higher macrophage levels at late time points compared to Pachόn. (Fig. 2h–i). This suggests that macrophages contribute to the prolonged immune response, potentially supporting debris clearance and cytokine production necessary for tissue repair.

Finally, B cells displayed a strikingly different pattern to neutrophils and macrophages. Both surface fish and Pachόn had minimal B cell numbers at early stages, but surface fish showed a robust increase after 7-dpci, increasing towards 14-dpci (Fig. 2e). RNAscope with pan-B cell marker *cd37*^+^ confirmed this late increase in B cells, with a higher density of B cells present in the surface fish compared to Pachón wounds (Fig. 2j), peaking at 30-dpci (Fig. 2k).

Our data suggest that the prolonged immune response in surface fish is driven by macrophages and B cells, with macrophages dominating the intermediate stages and B cells playing a major role in later stages. While macrophages are known to be important for phagocytosis, cytokine production, and collagen deposition during the early stages of fish heart regeneration, their roles in late-stage regeneration remain unexplored. Furthermore, the role of B cells in fish heart regeneration has not been characterised.

### Late-present surface fish immune cells exhibit distinct functional profiles compared to Pachόn

To investigate whether the immune cells with prolonged presence in surface fish have specific functions promoting regeneration at late stages of the injury response, we plotted the top five markers for macrophages, and B cells at 1-, 3-, 7-, 14-dpci, as well as in uninjured and sham controls in both fish (Fig. 3a–d). This revealed dynamic profiles over time in surface fish, with distinct gene sets activated at each time point, including late stages, indicating that the function of these cells changes over time with unique roles emerging during regeneration. In contrast, the transcriptional profiles of immune cells in Pachón largely returned to baseline by 7-dpci, reflecting a transient activation.

**Fig. 3.**
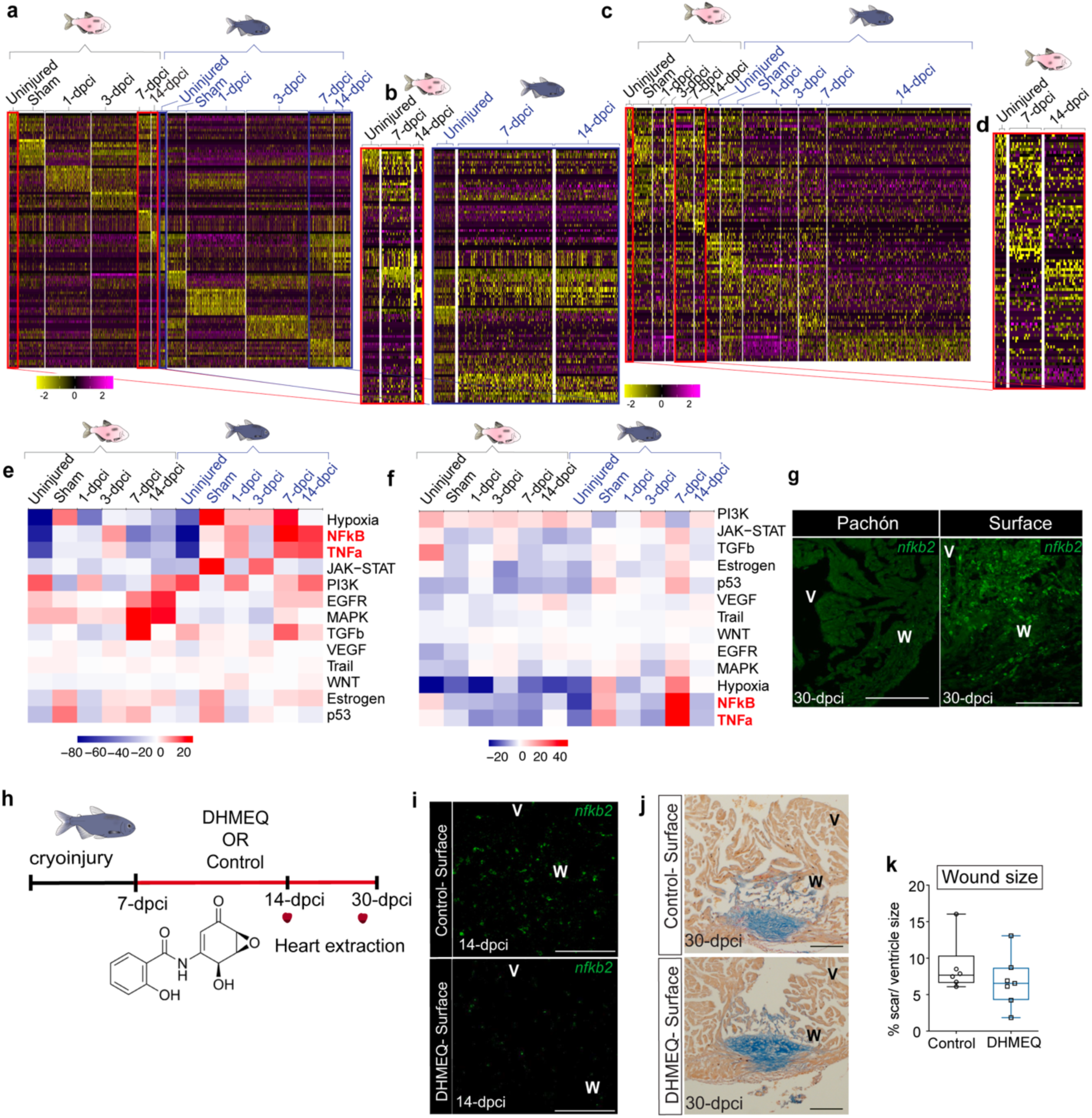
Distinct immune cell dynamics and pathway activation in surface fish and Pachόn at late stages after injury, with no direct impact of NF-κB inhibition on regeneration. a-c Heatmap displaying the top 5 markers for macrophages at all time points (a) and macrophages at 7- and 14-dpci zoomed in (b), and B cells at all timepoints in Pachón and surface fish (c) and B -cells at 7- and 14-dpci in Pachόn (d), respectively, shown across different time points: 1-, 3-, 7-, and 14-days post injury (dpci), as well as uninjured and sham controls. e-f PROGENy pathway analysis of marker genes shows upregulation of TNFα and NF-κB signalling pathways in surface fish macrophages (e) and B cells (f) at late stages. g, RNAscope to reveal *nfkb2*^+^ cells in the wound of Pachón and surface fish ventricles at 30-dpci, scale bar 100-μm all sections were counterstained with DAPI. h surface fish were injected with either Saline control or DHMEQ from 7- to 14-dpci and 7- to 30-dpci after cryoinjury, hearts were collected at 14-dpci and 30-dpci for RNAscope and AFOG staining, respectively. i RNAscope targeting *nfkb2* to confirm the effect of DHMEQ inhibition in the wound in Pachón and surface fish ventricles at 14-dpci shows decreased expression in DHMEQ-injected surface fish hearts, scale bar 100-μm, counterstained with DAPI. j AFOG-stained images of control and DHMEQ-injected surface fish ventricles at 30-dpci, scale bar 100-μm. k Quantification of the wound area in control and DHMEQ-injected hearts at 30-dpci shows no significant difference, p adjusted=0.61, unpaired t-test, n=6,7. DHMEQ, Dehydroxymethylepoxyquinomicin; dpci, days post cryo-injury; V, ventricle; w, wound.

To explore the pathways uniquely activated in the late-stage leukocytes, we performed PROGENy pathway analysis^26^ on macrophages and B cells. Despite the overall decline in macrophage numbers at later time points, this analysis revealed significant upregulation of key pathways such as TNFα and NF-κB signalling in both cell populations in surface fish at late stages (Fig. 3e-f). This persistent activation in the regenerating hearts suggests that even reduced quantities of these immune cells may contribute to the later stages of regeneration. The activation of TNFα and NF-κB contradicts their conventional roles as pro-inflammatory signals that typically peak during the early stages of injury and are thought to inhibit regeneration when prolonged^27–29^. This finding highlights the possibility that, in a regenerative context, these pathways may serve a dual function, initially driving inflammation but later contributing to tissue remodelling and repair.

To validate the potential pro-regenerative role of NF-κB signalling, we first examined the presence of *nfkb2*^+^ cells in the ventricles of both surface and Pachón fish at 14- and 30-dpci using RNAscope. Surface fish exhibited stronger NF-κB activation at both time points compared to Pachón (Fig. 3g). To further assess the functional impact of NF-κB at these late stages, we treated surface fish with Dehydroxymethylepoxyquinomicin (DHMEQ), an NF-κB inhibitor, from 7- to 14-dpci or 30-dpci (Fig. 3h). RNAscope analysis confirmed a significant reduction in *nfkb2*^+^ cells in DHMEQ-treated surface fish at 14-dpci compared to controls, indicating effective inhibition of NF-κB signalling (Fig. 3i). However, there was no observed reduction in B cells, macrophages, or *nfkb2*^+^ populations within these cell types (Supplementary Fig. 3a-e). Moreover, there was no significant difference in wound size between DHMEQ-treated and control surface fish hearts at 30-dpci (Fig. 3j–k). These results suggest that although late-stage NF-κB activation is prominent, its direct impact on cardiac regeneration in surface fish may be limited (Fig. 3k). While our scRNA-seq data shows that macrophages and B cells have unique functions at late stages, information from pathway analysis was limited.

### Late-stage phagocytic macrophages are required for cardiac regeneration

Since surface fish exhibit a prolonged presence of functionally distinct immune cells at late stages of healing, we further examined the functions of the macrophages during late stages of healing. Gene Ontology (GO) term analysis of the top cell markers for surface fish macrophages revealed enrichment of fibrosis-related terms (highlighted in red) at both 7- and 14-dpci, alongside pathways involved in immune activation, cytokine signalling, and tissue remodelling (Fig. 4a–b). Quantification of *c1qa*^+^ macrophages in surface fish hearts at 14-dpci following Dexamethasone treatment revealed no significant difference in the number of *c1qa*^+^ cells between treatment and control hearts (Fig. 4c). This observation suggests a complex interplay between macrophage persistence and inflammatory signaling during the later stages of regeneration. While NF-κB and other inflammatory pathways are upregulated, the maintenance of macrophage numbers appears to be regulated by additional factors beyond acute inflammatory signals. Given our GSEA analysis (Fig. 4a-b, Supplementary Fig. 4a–b) fits well with the well-established role of macrophages in phagocytosis and regulating fibrosis across multiple tissue contexts, we next investigated whether late-stage macrophages are actively phagocytic. Surface fish were injected with fluorescently labelled Dil PBS liposomes from 7- to 14-dpci, and hearts were harvested at 14-dpci. Approximately 75% of *c1qa*^+^ macrophages had ingested Dil liposomes (Fig. 4d, yellow arrows, Supplementary Fig. 4c), confirming that these macrophages are actively phagocytic at late stages of injury. To assess the functional importance of phagocytic macrophages, we depleted them using Clodronate liposomes from 7- to 12-dpci (Fig. 4e). Clodronate liposomes selectively deplete macrophages by inducing apoptosis in phagocytic cells upon uptake, allowing for targeted investigation of phagocytic cell function in tissue repair. Clodronate treatment significantly reduced the number of *c1qa*^+^ macrophages in the wound compared to PBS controls (Fig. 4f). At 30-dpci, Clodronate-treated hearts exhibited significantly larger wounds compared to the controls (Fig. 4g–h), indicating that late phagocytic macrophages with prolonged presence are essential for regeneration. In the absence of phagocytosis, overall cell proliferation in the wound was increased, which may explain the observed larger scars (Supplementary Fig. 4d). To further examine this, we specifically analysed proliferation in fibroblasts in Clodronate-treated surface fish hearts at 14-dpci (Fig. 4i). Clodronate-treated hearts exhibited a strong *tcf21*^+^ fibroblast proliferation, consistent with excessive fibrosis. In contrast, PBS-treated control hearts displayed fewer proliferating fibroblasts, suggesting that macrophages help restrain fibroblast expansion and limit scarring during regeneration, consistent with previous results in the salamander^11^. These findings indicate that the macrophages present in later stages mainly function through phagocytosis to reduce scarring.

**Fig. 4.**
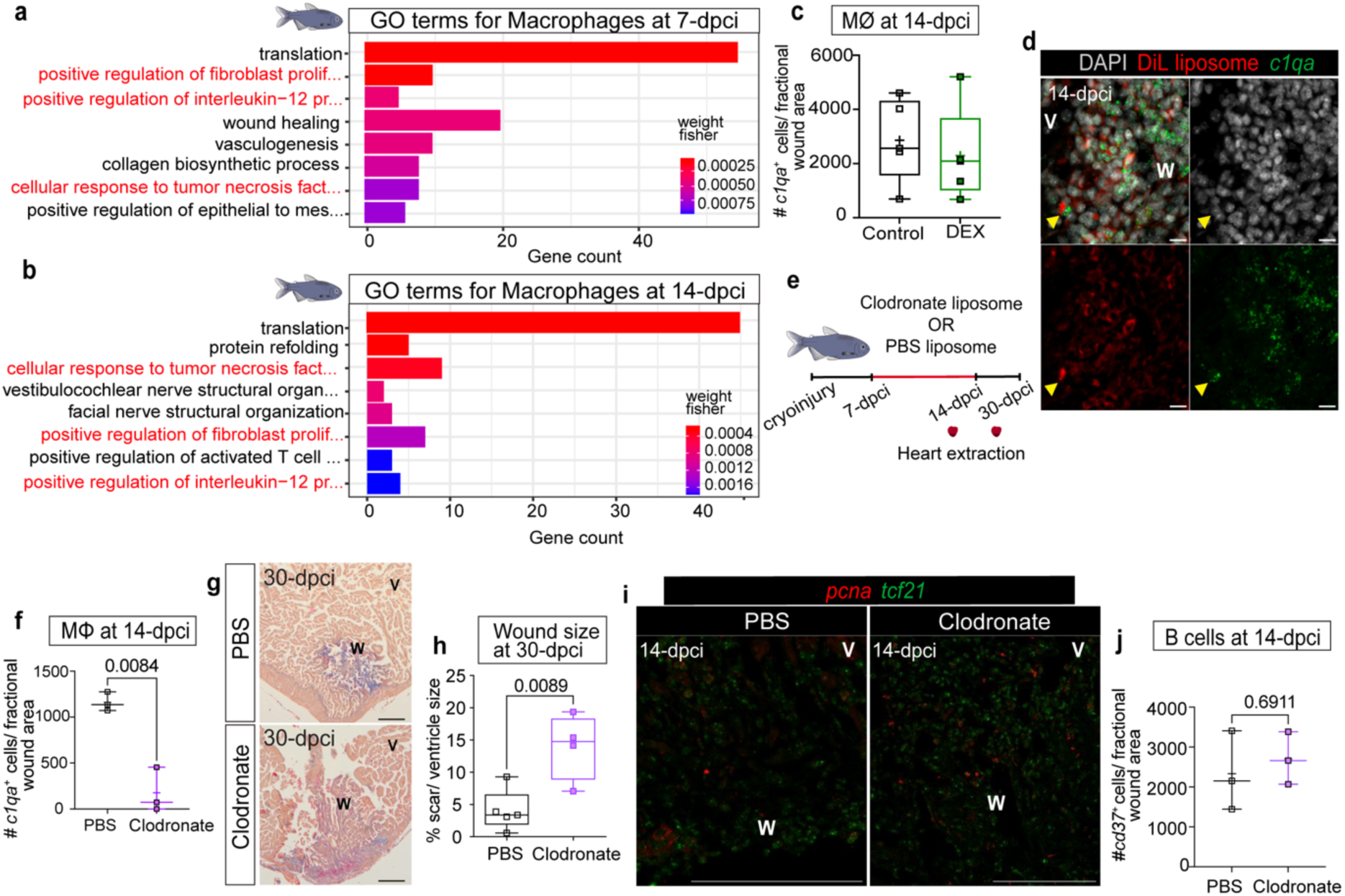
surface fish late-macrophages are functionally active and essential for regeneration. a-b. GO term analysis for top cell markers of surface fish macrophages at 7-dpci (**a**) and 14-dpci (**b**). GO terms related to fibrosis are highlighted in red. **c** Quantification of RNAscope targeting *c1qa^+^* cells at 14dpci in Dex-treated and control surface fish hearts, unpaired -t-test, n=4/ each group. **d** surface fish were injected with fluorescently labelled DiL PBS liposomes from 7- to 11-dpci after cryoinjury, hearts were collected at 14-dpci, confocal image of *c1qa* (green), fluorescent PBS liposomes (red), and DAPI (grey) co-localisation patterns in the heart, yellow arrow points macrophage ingested fluorescent PBS liposome (75% of macrophages colocalised with DiL liposome, confidence Interval: 58.80% to 90.35% **e** Surface fish were injected with PBS (as control) and Clodronate liposomes from 7- to 11-dpci after cryoinjury, hearts were collected at 14-dpci for RNAscope and Mef2/PCNA immunostaining and at 30-dpci and AFOG staining. **f** Quantification of *c1qa*^+^ cells present in the wound at 14-dpci in PBS vs Clodronate treated surface fish hearts show macrophages were significantly depleted upon Clodronate liposome injection, unpaired t-test, n=3/ each group. **g** AFOG-stained images PBS-liposome as control or Clodronate-liposome injected surface fish ventricles at 30-dpci, scale bar 100-μm. **h** Quantification of the difference in wound area between PBS-liposome control or Clodronate-liposome treated hearts at 30-dpci shows Clodronate treatment significantly impaired regeneration in surface fish, unpaired t-test, n=4-5. **i** Representative confocal image of fibroblast marker *tcf21* (green), proliferation marker *pcna* (red), and nuclear DAPI (grey in the IP-PBS control and IP-Clodronate surface fish hearts at 14-dpci. **j** Quantification of *cd37*^+^ cells present in the wound at 14-dpci in PBS vs Clodronate treated surface fish hearts shows no difference in B cell numbers when surface fish injected with Clodronate, unpaired t-test, n=3 each group dpci, days post cryo-injury; V, ventricle; W, wound.

Since macrophages are known to play a central role in modulating the immune response, we examined whether their depletion would impact other immune cell populations in the wound, such as B cells. Clodronate-treated surface fish hearts showed no reduction in B cell numbers compared to PBS-treated controls (Fig.4j) suggesting that macrophages are not important for recruitment of B cells. Additionally, as B cells can exhibit phagocytic activity in some contexts^30–32^, we only observed colocalisation specifically with *c1qa*-expressing cells, further supporting the idea that macrophages are the primary phagocytes in this context. If B cells contribute to regeneration, it is likely through mechanisms other than phagocytosis (Fig. 4d). Furthermore, when we quantified the number of B cells recruited into the wound after injury in the Panther zebrafish mutant, which lacks *csf1ra* macrophage populations, we did not observe any difference in B cell numbers between Panther and the control zebrafish (Supplementary Fig. 4e–f). Together, this shows that B cell recruitment or activation during cardiac repair is independent of macrophage presence.

### Late-present B cells may have a leading role in cell cycle control and proliferation during heart regeneration

Following the demonstration that late-stage macrophages are critical for cardiac regeneration through their phagocytic activity, we sought to explore the role of B cells. To investigate the observed contribution of B cells to regeneration (Fig. 1h–i), we analysed *cd37*^+^ B cells in surface fish hearts at 14- and 30-dpci following Dexamethasone treatment. We observed a significant reduction in *cd37*^+^ cell numbers at both 14- and 30-dpci compared with controls (Fig. 5a–b), indicating that, in contrast to macrophages, B cell activity is impaired by Dexamethasone and that recruitment of B cells relies on pro-inflammatory signals.

**Fig. 5.**
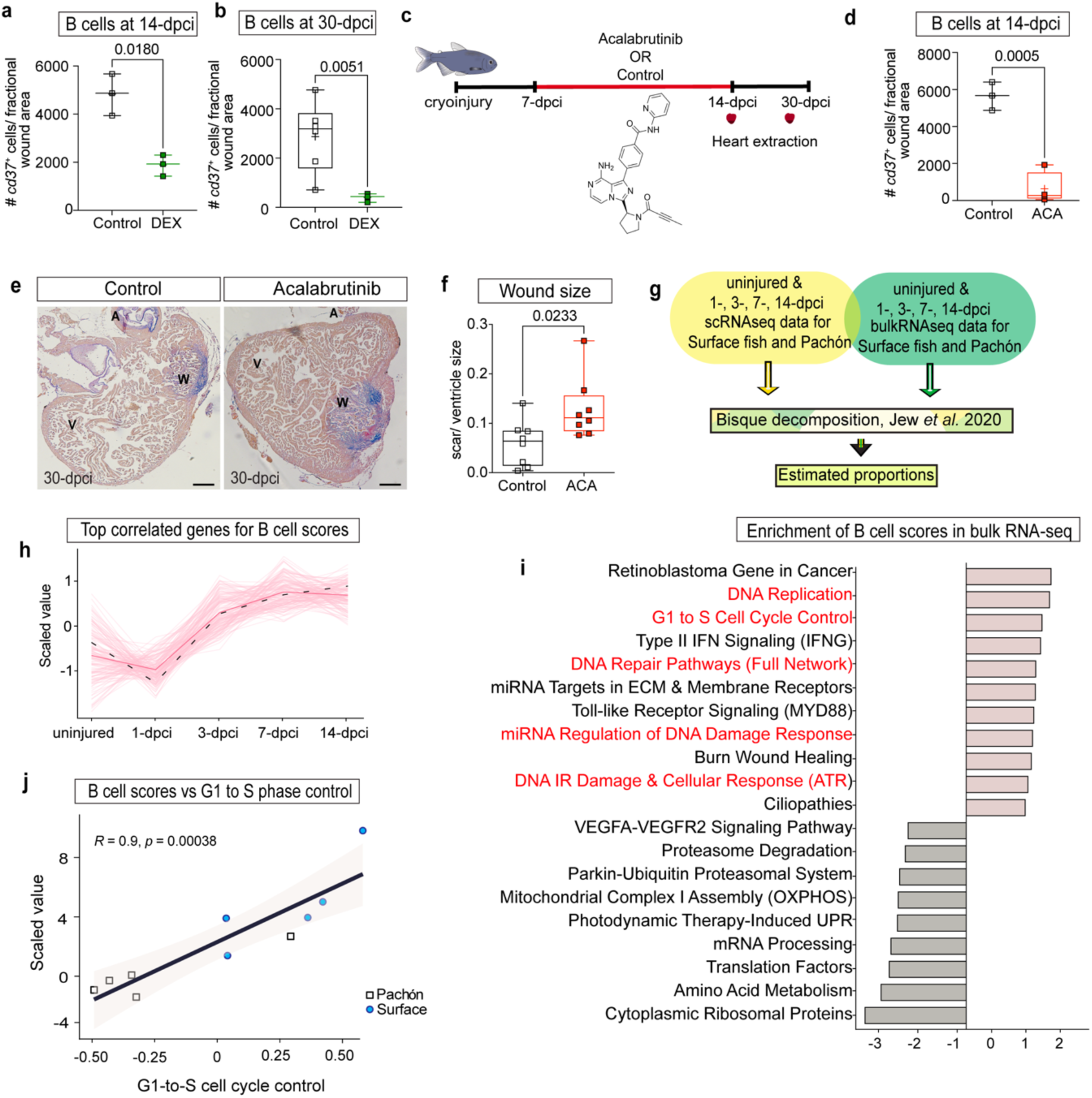
Temporal dynamics of B cells and their involvement in cell cycle regulation during cardiac regeneration in surface and Pachón fish. a Quantification of RNAscope targeting *cd37^+^* cells at 14-dpci, b and at 30-dpci in Dexamethasone-treated and control surface fish hearts show significantly impaired B cell response upon Dexamethasone treatment in both time points, unpaired t-test, n=3/each group and n=5, 3, respectively. c surface fish were injected with either control (corn oil, DMSO) or Acalabrutinib from 7- to 11-dpci, hearts were collected at 14-dpci and 30-dpci for RNAscope, immunostainings and AFOG staining. d Quantification of *cd37*^+^ cells present in the wound at 14-dpci in control vs Acalabrutinib-injected surface fish hearts shows B cells were significantly depleted upon inhibitor treatment, n=3, 4. e AFOG-stained images of control and Acalabrutinib-injected surface fish ventricles at 30-dpci, scale bar 100-μm. f Quantification of the difference in wound area between the control and Acalabrutinib-injected surface fish hearts at 30-dpci shows significantly larger wounds in Acalabrutinib group compared to control, unpaired t-test, n=7, 8. g Immune deconvolution workflow using single-cell RNAseq and bulk RNAseq data for surface fish and Pachόn at 1-, 3-, 7-, and14-dpci, and uninjured. h Temporal dynamics of B cell activity show increased B cell scores in surface fish starting at 7-dpci, with lower activity at earlier time points. i WikiPath Pathway enrichment analysis in the bulk RNAseq data highlights significant upregulation of proliferation, cell cycle regulation, and wound healing pathways in surface fish B cells. j Correlation between B cell scores and G1-to-S phase cell cycle control pathway shows a correlation in surface fish, indicating that B cell activity is associated with cell cycle regulation. In contrast, Pachón fish exhibit a negative correlation, suggesting impaired regeneration. DEX, Dexamethasone; ACA, Acalabrutinib; dpci, days post injury; NES, Normalised Enrichment Score.

To dissect the role of B cells, surface fish were treated with Acalabrutinib, a Bruton tyrosine kinase (BTK) inhibitor that specifically targets B cells, from 7- to 11-dpci, and hearts were analysed at 14- and 30-dpci (Fig. 5c). Acalabrutinib treatment significantly reduced *cd37*^+^ B cell numbers in the wound area at 14-dpci compared to controls (Fig. 5d). At 30-dpci, Acalabrutinib-treated hearts exhibited significantly larger wounds compared to controls demonstrating that B cell depletion impairs regeneration (Fig. 5e–f).

Single-cell analysis of B cells was challenging due to their low number in Pachón cavefish and the high proportion of functionally unannotated genes in our dataset, making it difficult to investigate their functions further using this data. To enable more comprehensive immune profiling of B cells, we used deconvolution to leverage our scRNA-seq data to estimate cell-type composition and abundance in bulk RNA-sequencing, which provides greater depth and sample size^33^. We performed immune deconvolution analysis on bulk RNA-seq data from Pachón and surface fish hearts at multiple time points (1-, 3-, 7-, and 14-dpci, and uninjured; data from Lekkos et al., unpublished) (Fig. 5g). We first validated our differential proportion analysis for the major immune cell types and observed the expected early immune response dynamics. Neutrophils showed a rapid induction, peaking at 1-dpci, followed by a peak in macrophages at 3-dpci and a prolonged elevated response, consistent with their trends observed in the earlier DPA results (Fig. 2, Supplementary Fig. 5a–b).

Consistent with our previous results, deconvoluted B cell scores were significantly elevated in surface fish compared with Pachón at 14-dpci (Supplementary Fig. 5c). To elucidate the processes associated with B cell activity in surface fish, we identified genes that were correlated with B cell scores across the injury time course, which increased at 7-dpci and were sustained at 14-dpci (Fig. 5h). Several pathways involved in proliferation, regulation of the cell cycle, and wound healing were significantly enriched in the positively correlated genes (Fig. 5i). Notably, deconvoluted B cell scores correlated robustly with GSVA summary scores for the G1-to-S phase cell cycle control pathway in surface and Pachón cavefish samples at 14-dpci (Fig. 5j). Therefore, B cells appear to be closely associated with overall proliferation rates in the heart. While it is known that cardiomyocyte proliferation is a critical driver of successful cardiac regeneration^1,10^, it has remained unclear whether B cells themselves could proliferate. To determine whether the observed correlation was due to B cell proliferation, we examined B cell-specific cell cycle activity. Analysis of scRNA-seq data for *pcna* expression in B cells revealed no significant activation of cell cycle markers in B cells from 7-dpci onwards (Supplementary Fig. 5d). As the B cells did not proliferate, the correlation between deconvoluted B cell scores and cell cycle pathways suggests that B cells may facilitate cardiomyocyte proliferation.

### B cells influence border zone cardiomyocyte proliferation after injury

Given that cardiomyocytes are the most abundant cell type in the heart, our bulk data may reflect a relationship between B cell activity and cardiomyocyte proliferation. Therefore, we next examined the potential role of B cells in regulating border zone cardiomyocyte proliferation. We have previously shown that cardiomyocyte proliferation was induced in both Pachón and surface fish after resection injury^19^. In contrast, surface fish exhibited robust cardiomyocyte proliferation with complete tissue repair. To investigate this further after cryoinjury, we assessed border zone cardiomyocyte behaviour in surface fish and Pachón before and after cryoinjury by subsetting the scRNA-seq data to isolate the cardiomyocytes based on *myl7* expression (Supplementary Fig. 6a). Re-clustering revealed all expected cardiomyocyte populations present during homeostasis, including cortical, trabecular, primordial populations, as well as the emergence of injury-responsive border zone cardiomyocytes after cryoinjury (Fig. 6a–b).

**Fig. 6.**
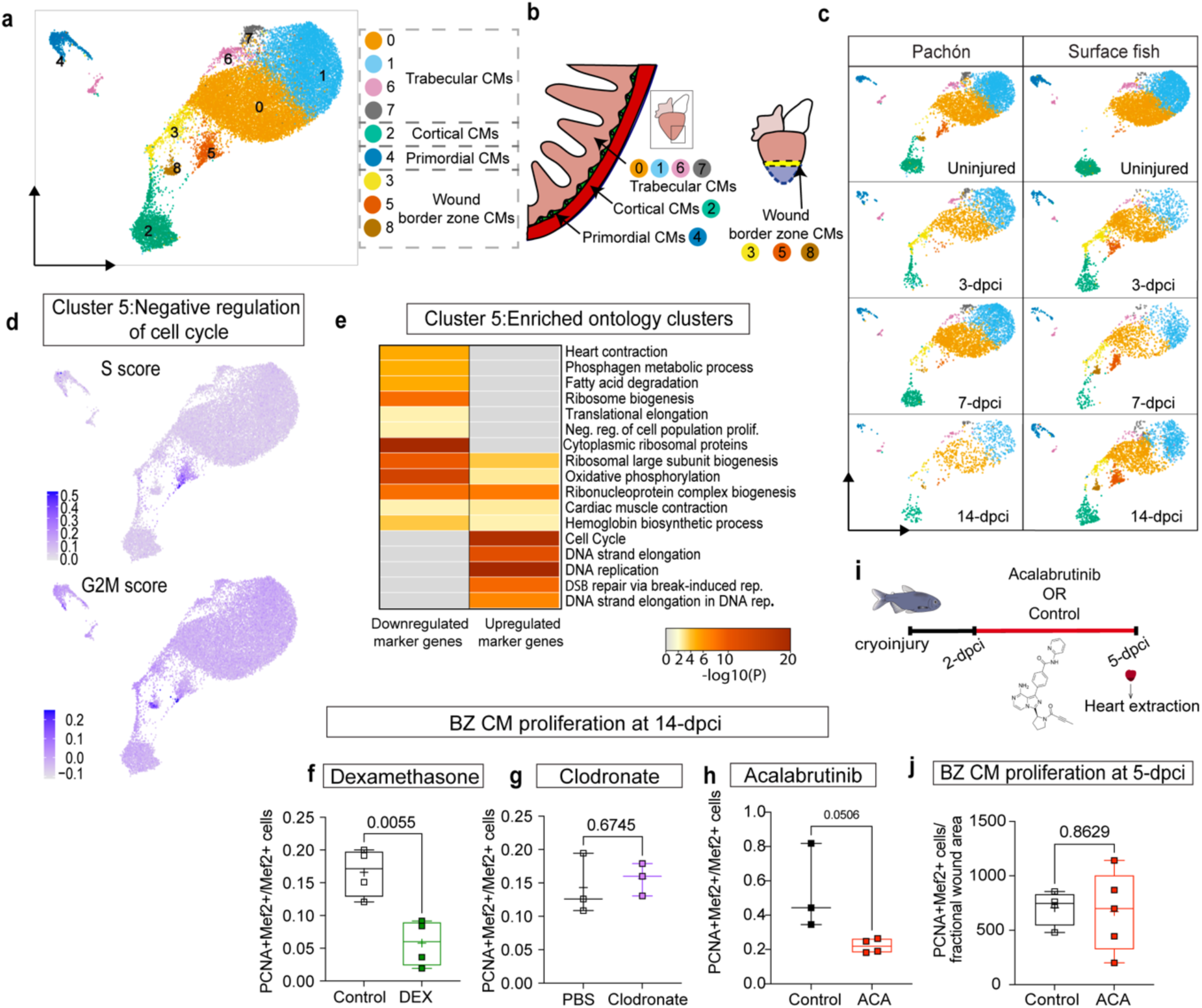
Border zone cardiomyocyte proliferation is disrupted in Pachόn due to negative regulation of the cell cycle. **a** Cardiomyocytes re-clustered and all the subsets labelled according to their marker gene expressions in *A. mexicanus* ventricles **b**. A schematic illustrating the diverse cardiomyocyte sub-populations within the heart: trabecular cardiomyocytes (0, 1, 6, 7), cortical cardiomyocytes (2), primordial cardiomyocytes (4), and wound border zone cardiomyocytes on the right plane (3, 5, 8). **c** UMAP showing all border zone cardiomyocyte clusters (cluster number 3, 5, and 8) are proliferating in surface fish after cryoinjury whereas although they are present un uninjured Pachόn they decrease after injury. **d** UMAPs showing cell cycle phases in Pachόn cardiomyocytes, with the cluster 5 border zone cardiomyocytes accumulating in S-phase. **e** Gene ontology terms for cluster 5 shows strong upregulation in Cell cycle and DNA repair pathways. **f-h** Quantification of Mef2 and PCNA double-positive cells show no difference between Dex-treated and control, n=4/ each group (**f**), Clodronate and PBS control n=3/ each group (**g**) and Acalabrutinib and control, n= 3, 4 (**h**) in proliferating cardiomyocytes at the border zone at 14-dpci. **i** Surface fish were injected with either control (Saline, corn oil, DMSO) or Acalabrutinib from 2- to 5-dpci, hearts were collected at 5-dpci for immunostaining. **j** Quantification of Mef2 and PCNA double-positive cells show no difference between the control vs Acalabrutinib-injected fish for the proliferating cardiomyocytes at the border zone, unpaired t-test, n=4, 5. DEX, Dexamethasone; ACA, Acalabrutinib; dpci, days post injury; BZ CM, border zone cardiomyocytes.

In both surface fish and Pachón, similar new clusters of cardiomyocytes appeared after injury. Border zone clusters 3 and 5 emerged at 3-dpci. Cluster 5 exhibited S-phase activity, indicating the initiation of DNA replication (Fig. 6c–d). GO analysis of cluster 5 revealed a significant upregulation of pathways related to the cell cycle and DNA repair (Fig. 6e), underscoring its involvement in DNA replication and initiation. By 7-dpci, M-phase cluster 8 appeared, indicating progression through the cell cycle (Supplementary Fig. 6b–c) demonstrating strong enrichment in chromosome and cell division processes, RNA processing, and protein and peptide metabolism, consistent with its role in mitotic progression (Supplementary Fig. 6d–e). In surface fish, all three clusters were present, suggesting that DNA replication was activated and followed by mitosis. In contrast, while cluster 3 was activated in Pachón, very few cardiomyocytes activated S-phase genes in cluster 5, and cluster 8 was almost absent following injury (Fig. 6c). Interestingly, these clusters were present in the uninjured Pachόn hearts, suggesting that cardiomyocyte proliferation in Pachón was higher before injury but becomes disrupted post-injury (Fig. 6c). The accumulation of Pachόn cluster 3 cardiomyocytes at 3- and 7-dpci but without subsequent mitosis aligns well with previous findings showing upregulation of *pcna* in the border zone cardiomyocytes, as *pcna* is expressed in G1 and S phase^34^. The appearance of M-phase cluster 8 coincides with the start of B cell recruitment in surface fish, further suggesting a correlation between the prolonged immune response and cardiomyocyte proliferation.

To understand if late cardiomyocyte proliferation was indeed regulated by the prolonged immune response, we first analysed border zone cardiomyocyte proliferation in sections of surface fish hearts treated with Dexamethasone or Clodronate. For Dexamethasone-treated surface fish hearts, myocardial proliferation was significantly decreased compared to controls at 14-dpci (Fig. 6f), whilst Clodronate-treated hearts showed no difference in the proliferation of border zone cardiomyocytes compared to controls at 14-dpci (Fig. 6g). This suggests that the prolonged presence of macrophages does not regulate proliferation, aligning well with our finding that they mainly have a phagocytotic function. In contrast, B cell depletion using Acalabrutinib resulted in a marked reduction of proliferating cardiomyocytes at the wound border at 14-dpci (Fig. 6h), indicating that B cells are indeed important for cardiomyocyte proliferation. To exclude the possibility of off-target effects of the inhibitor treatment directly affecting cardiomyocyte proliferation, we injected fish with Acalubritinib at 2-, 3-, 4-dpci and isolated the hearts at 5-dpci, when proliferation was already initiated but before B cell recruitment occurs in surface fish (Fig. 6i). We observed no difference in cardiomyocyte proliferation at 5-dpci (Fig. 6j), confirming that the B cell response is critical for supporting cardiomyocyte proliferation and efficient wound healing during cardiac regeneration.

### Role of B cells in regeneration across species

To further validate our findings in a different species, we turned to the zebrafish model, which shares many regenerative features with surface fish. We analysed IgM^+^ cell populations in adult zebrafish hearts following cryoinjury (Fig. 7a). IgM is the major immunoglobulin and a well-established marker for the predominant subset of B cells in zebrafish. Quantification revealed a significant increase in IgM^+^ cells in in Tg(*Cau.ighv-ighm:EGFP*)^35^ hearts at 14- and 21-dpci compared to unwounded controls (Fig. 7b–c). To test the functional role of B cells in zebrafish regeneration, we used the *rag1^-/-^* mutant, which lacks mature lymphocytes, including B cells^36,37^. At 21- and 60-dpci, wild-type hearts exhibited robust regeneration, while *rag1^-/-^* hearts showed impaired repair, as evidenced by quantification of wound size at 21- and 60-dpci (Fig. 7d–e). Similar to the observations in surface fish, we found a significant reduction in proliferating cardiomyocytes at the border zone in *rag1^-/-^* hearts compared to wild-type controls (Fig. 7f).

**Fig. 7.**
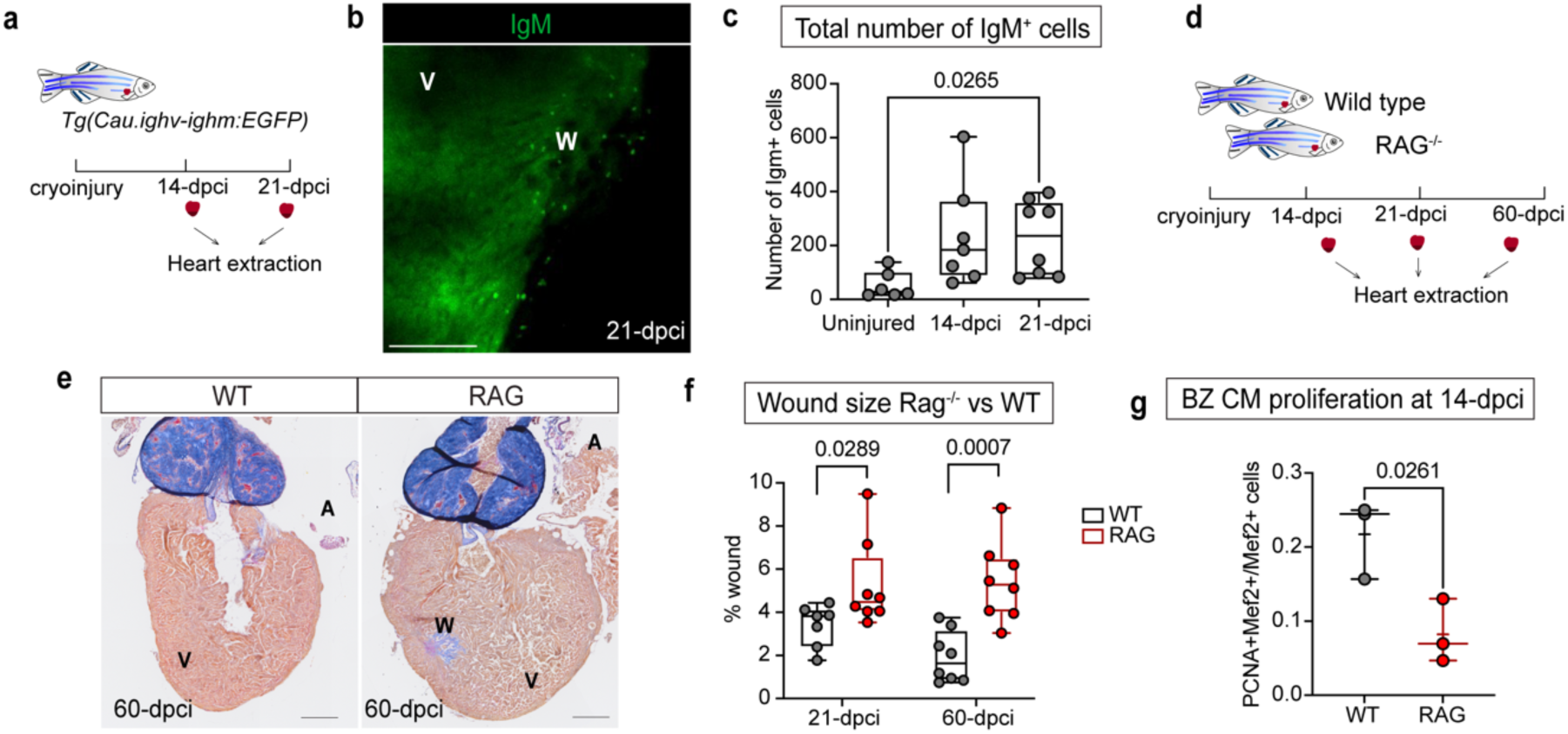
Cross-species validation of B cell involvement in cardiac regeneration. **a** Schematic indicating the timeline of zebrafish IgM experiments. **b** Representative image of 21-dpci Tg(*Cau.ighv-ighm:EGFP*)^35^ zebrafish heart showing IgM cells in the wound zone, scale bar 100-μm. **c** Quantification of IgM^+^ cells in Tg(*Cau.ighv-ighm:EGFP*) hearts at 14- and 21-dpci as well as unwounded controls reveals a significantly elevated IgM response at 21-dpci, n=6-8/ each group, unpaired t-test. **d** Wild Type and *rag1*^-/-^ zebrafish were cryoinjured and the hearts were collected at 14-, 21- and 60-dpci for RNAscope, immunostainings and AFOG staining. **e** Representative images of wild-type and *rag1*^-/-^ zebrafish hearts at 60-dpci, the wound area is indicated with black dashed line, scale bar 100-μm. **f** Quantification of wound size of *rag1*^-/-^ and wild-type control hearts at 21- and 60-dpci shows significantly larger wounds in *rag1*^-/-^ mutant hearts, n=6-8/ each group, Two-way ANOVA. **g** Quantification of Mef2/ PCNA double-positive cells show significant difference between wild-type control vs *rag1*^-/-^ zebrafish for the proliferating cardiomyocytes at the border zone, unpaired t-test, n=3/ each group. W: wound, A: atrium, V: ventricle; WT, wild type, dpci, days-post-injury.

These findings indicate that the B cell response is conserved among different regenerative species and plays a critical role in promoting cardiomyocyte proliferation and efficient cardiac regeneration.

## Discussion

In this study, we leveraged scRNA-seq and immune profiling to identify key factors influencing the difference in cardiac regeneration between surface and Pachón subtypes of *A. mexicanus* and further verified our B cell findings in regenerative adult zebrafish. We observed a significantly diverging immune response between regenerative surface fish and scarring Pachón following cryoinjury. Surface fish exhibited a prolonged activation of macrophages and B cells, which were associated with successful cardiac repair, while Pachón showed a more transient immune response. These findings, showing that a prolonged immune response is crucial for successful regeneration, are surprising as extended inflammation is considered detrimental to regeneration.

Studies involving macrophage ablation in zebrafish and neonatal mice have focused on the early days after injury, showing that disrupting the initial macrophage influx inhibits regeneration^9,13,25,38^. Similarly, Medaka, another teleost fish species that exhibits limited cardiac regeneration unlike zebrafish, completely lack the macrophage response. While Pachόn also do not regenerate, their response differs significantly from Medaka, indicating key differences between fish species. Pachόn exhibit a remarkably similar initial influx of neutrophils and macrophages to regenerative surface fish, yet regeneration is still unsuccessful. It is only after the initial peak that significant differences which are crucial for regeneration appear in the immune response between the two morphs, which are crucial for regeneration. We were able to pick up on these more subtle differences as *Astyanax mexicanus* provides the unique opportunity to directly compare between two morphs of the same species with different regenerative abilities, surface fish and Pachόn. This provides the granularity needed to identify fine nuances in the response that determine the difference between successful regeneration and scarring, which might be lost when comparing entirely different species or using blunt cell ablation approaches.

The beneficial role for a prolonged presence of macrophages during regeneration is surprising, especially as these cells upregulate pro-inflammatory genes such as *tnfa* and *nfkb2* specifically at these late stages. The prevailing consensus in the field is that a balance between pro-inflammatory and reparative gene expression in macrophages is key to regeneration and that prolonged pro-inflammatory activation of macrophages disrupts regeneration^39,40^. However, our findings indicate that while the late upregulation of pro-inflammatory genes does not positively contribute to regeneration, it also does not hinder it, as would be predicted based on previous studies^9,13^. NF-κB has been shown to be required for cardiomyocyte proliferation during regeneration^41^ but we did not find an effect on regeneration when inhibiting NF-κB during late stages, indicating the effect of NF-κB on cardiomyocyte proliferation occurs during the earlier stages of regeneration. Prolonged NF-κB and TNFα signatures might be key to maintaining specific cellular functions of macrophages throughout late stages of regeneration. For example, TNFα has been shown to enhance phagocytosis in macrophages by upregulating phagocytic receptors and stimulating reactive oxygen production^42,43^. Tissue-specific targeting of NF-κB and TNFα will be required to better understand their role at these late stages, as overall inhibition did not show effects on regeneration. Phagocytosis, a key task of macrophages during early stages^9,25,44^, indeed appeared to be a crucial function of late-stage macrophages. Most macrophages present after 7-dpci were actively phagocytic, suggesting a role in wound clearance and tissue remodelling. It is only after 7-dpci that the wound strongly starts to reduce in size, but the mechanisms of wound clearance have not yet been established. Ablation of phagocytic macrophages using Clodronate liposomes also enhanced proliferation of fibroblasts in the wound, suggesting that lack of clearance of fibroblasts allows these cells to settle and initiate proliferation to establish a permanent scar. Although this could also be caused by direct signalling^11,45,46^.

While our scRNA-seq data suggested a prolonged initial peak in neutrophil influx in Pachón compared to surface fish, as well as low-level retention in surface fish after 3-dpci compared to Pachόn, neither observation was confirmed in tissue sections. Neutrophils are known to have dual roles in cardiac injury. A prolonged initial influx, as seen in the single-cell data in Pachón, has been strongly linked to scarring in the mammalian heart^47^, while certain pro-regenerative neutrophil states have also been described^48–50^. Although neutrophils may not be the primary drivers of the differences in regeneration between surface fish and Pachόn, their role in the early inflammatory response and potential contribution to tissue repair need further elucidation.

In contrast to neutrophils, we confirmed B cells as a critical cell type involved in late-stage regeneration. Large numbers of B cells were recruited in both surface fish and zebrafish during late stages of regeneration. Inhibition of B cells using Acalabrutinib in surface fish or through mutation of *rag1* in zebrafish resulted in reduced cardiomyocyte proliferation and regeneration, indicating the positive influence of B cells on regeneration is conserved between species. While RAG mutants lack mature B and T cells due to defective V(D)J recombination, a critical process for adaptive cell development^37,51^, our data suggest that the effect on regeneration is due to B cell loss, as T cells are not prominently involved in late stages of cardiac regeneration and their numbers remain low. Our findings fit well with recent findings from neonatal mice^17^, in which depletion of B cells was found to reduce cardiomyocyte proliferation and impair cardiac function. Regulatory B cells (Bregs) have also been shown to improve ventricular remodelling after MI by modulating monocyte migration and reducing inflammation^52^. Secretion of IL-10 by Bregs creates a pro-regenerative environment that may indirectly support cardiomyocyte proliferation by reducing inflammation and fibrosis, with a similar function found for bone marrow-derived naïve B cells^53^. This suggests that B cells play a conserved role in supporting cardiac repair across different injury models, although we could not find conserved mechanisms, as IL-10, for example, was not highly expressed in surface fish B cells. In non-mammalian models, B cells have not yet been implicated in tissue regeneration, and it has long been suggested that there is an inverse relationship between levels of adaptive immune response and ability to regenerate^11,54^. However, our findings indicate that a well-established adaptive B cell response, particularly in surface fish, supports the regenerative process. The effect of B cells on cardiomyocyte proliferation is somewhat unexpected as proliferation is initiated from 3-dpci, while B cell invasion only takes place from 7-dpci onwards. As border zone cardiomyocytes progress from S- to M-phase between 3- and 7-dpci, B cells could play a role in promoting cell cycle progression into mitosis. Mammalian B cells secrete IL-10, a cytokine that has been shown to specifically affect cell cycle progression into M-phase in tendon-derived stem cells, mediated through the JAK/Stat3 pathway^55^. A similar mechanism could apply to fish cardiomyocytes, where cytokines secreted by B cells may drive cell cycle progression into mitosis and subsequent cytokinesis, an essential part of successful regeneration, which does not occur in their mammalian equivalents^56^. While we focus on the role of B cells during cardiomyocyte proliferation, in mammalian models, B cells have also been shown to modulate regeneration through tissue remodelling and angiogenesis. For example, in mice, B cells have been shown to regulate fibrosis and inflammation through the production of cytokines, which in turn influence macrophage polarisation and extracellular matrix remodelling^57,58^. Comparison of the functions of surface fish and Pachón B cells in the single-cell RNA-seq data did not provide much information on other functions due to the very low number of these cells in Pachón, combined with limitations in gene annotation. Future work will need to explore the molecular interactions between B cells and the other immune cell populations, as well as their role in modulating cardiomyocyte proliferation, the extracellular matrix and promoting angiogenesis during cardiac repair.

The surface fish were found to have a prolonged innate and adaptive response, with not only the neutrophils and macrophages, but also B cells present at all time points, including late stages. As this immune cell profile was absent in Pachón, which has a strongly altered metabolism^59,60^, this raises the possibility that immune cell populations in Pachón do not have the energy to sustain a long-term response as a result of a metabolic trade-off favouring survival over costly immune functions. In resource-limited environments, species often prioritise innate immunity over adaptive immunity due to lower energetic costs of innate defences, as seen in birds and rodents (e.g., B cell-mediated responses)^61,62^. In case of Pachόn, the loss of sustained B cell activity and other immune functions may reflect a similar trade-off, where adaptation to nutrient poor cave environment comes at the expense of adaptive immune responses. In turn, this suggests that surface fish and zebrafish have unique metabolic advantages over non-regenerative species that allow leukocytes to survive long-term and complete the regenerative process^63,64^.

In conclusion, our study demonstrates that a prolonged immune response, driven by sustained activation of macrophages and B cells, is critical for successful cardiac regeneration. The transient immune response in Pachόn is characterised by insufficient immune activity and lack of a B cell response. These findings provide a framework for understanding the complex interplay between immune cells and tissue repair, with potential implications for improving regenerative strategies in mammals, including humans.

### Animal Husbandry

All *Astyanax mexicanus* procedures were conducted at the aquatic animal facilities of Biomedical Services and the Institute of Developmental and Regenerative Medicine at the University of Oxford. These procedures comply with the revised UK Animals (Scientific Procedures) Act 1986, Directive 2010/63/EU of the European Parliament and the Council of 22 September 2010 on the protection of animals used for scientific purposes, Home Office licenses, and were approved by Oxford University’s central Committee on Animal Care and Ethical Review (ACER).

All *Astyanax mexicanus* experiments were performed using wild-type surface fish from the Río Choy background and cavefish from the Pachón cave background. The fish were maintained in system water at 21–23°C under a 14:10-hour light/dark cycle. *Danio rerio* (zebrafish) experiments were conducted at the University of Bristol in accordance with UK Home Office and by the local University of Bristol Animal Welfare and Ethical Review Body (AWERB). The Tg(*Cau.ighv-ighm:EGFP*)^35^ and *rag1^hu^*^1999^ ^36^ mutants have been described previously and were maintained under standard conditions. Age matched wild-type fish served as controls for all *rag1*^-/-^ experiments.

### Cardiac Injury

Cryoinjury was performed as the preferred method of cardiac injury, following a previously described protocol for zebrafish^65^. Briefly, fish were anesthetised in 0.25 g/L MS222 and transferred to a wet sponge. Scales on the thorax were removed with forceps, and a small incision was made using spring scissors. The pericardium was carefully removed, and the heart surface was dried using sterile lint-free tissue. Cryoinjury in *A. mexicanus* was performed inside the chest cavity, whereas in zebrafish, it was applied to the surface of the incision as the heart elevates when slight pressure is applied to the abdomen. A copper cryoprobe (0.5 cm diameter) was cooled in liquid nitrogen for a few seconds and placed on the ventricle until thawed. Injury size was monitored by observing colour changes in the area, and recovery was completed within 1–2 minutes by pipetting water over the gills. surface fish were individually housed post-surgery to minimise stress responses. For drug treatments, fish were randomly assigned to treatment groups at the end of the operation. Sham surgeries involved making a thorax incision, removing the pericardial sac, and placing a room-temperature copper probe on the ventricle before returning the fish to system water. All procedures were conducted at the same time of day by the same individual to minimise variability. Control hearts were isolated from uninjured fish, while sham group hearts were isolated 3 days post-operation. Cryoinjured hearts were isolated at various time points between 1 and 60 days for cryoinjury.

### Single-cell RNA sequencing and analysis

Ventricles were isolated, digested, and processed into a single-cell suspension according to the digestion method presented in Potts et al, 2022^66^. Briefly, ventricles were digested using a buffer containing PBS, taurine, glucose, 2,3-butanedione monoxime, HEPES, CaCl₂, collagenase II/IV, and DNase I. Tissue was digested at 32°C, and dissociated cells were filtered using 100-μm filters before resuspension in DMEM with 10% foetal bovine serum.

Single-cell RNA sequencing was performed using the 10X Chromium Next GEM Single Cell 3’ v3.1 kit. Sequencing was conducted on the Illumina NextSeq 500/550, achieving ∼50,000 reads per cell. Fastq files were generated using Cell Ranger mkfastq (v3.0.2), and reads were aligned to *A. mexicanus* genome assembly (v1.0.2) with custom 3’UTR-extended annotations^66^

Analysis was conducted using Seurat (v4.0.6)^67^. Quality control removed low-quality cells based on average nFeatures and nCounts values and data were normalised using SCTransform^68^. Batch effects were corrected for during integration of all 12 samples using Harmony (v1.0)^69^ to create a final data of 85,516 cells. The integrated dataset was centred and scaled prior to dimensional reduction using PCA and UMAP. Appropriate cell cluster resolution was set using Clustree^70^ and cell clusters were found, identifying all major cardiac cell types expected in the heart before and after injury. Cell types were annotated based on marker genes. Differential proportion analysis was performed according to the methodology presented in Farbehi et al^71^. Differential gene expression was analysed using MAST^72^, Gene ontology^73^ and pathway analysis were performed using fGSEA (v1.16.0)^74^, and PROGENy (v1.12.0)^26^.

### Drug Administration

Water-soluble Dexamethasone was administered at a concentration of 10mg/L in system water for 7 consecutive days. The Dexamethasone solution and control system water were replenished daily for both the treatment and control groups.

Clodronate liposomes (5mg clodronate/mL) and control PBS liposomes (Liposoma, the Netherlands) were administered via intraperitoneal injections between 7– and 11-dpci. DiL fluorescent PBS liposomes were injected from 7– to 11dpci for five consecutive days to visualise affected cell populations. Hearts were isolated at 14- and 30-dpci for RNAscope and immunostaining.

Acalabrutinib was administered intraperitoneally between 7- and 11-dpci, and hearts were isolated at 14- and 30-dpci. The concentration was determined as 5 mg/kg based on murine studies. A working solution was prepared by diluting a 10 mg stock solution (1:10 DMSO: corn oil) to 0.5 mg with corn oil. The injection solution was sonicated in a Bioruptor Pico ultrasonicator using the following program: 30 seconds of sonication at ultrahigh frequency,followed by a 30-second break, repeated for 30 cycles at 20°C. Fresh injection solutions were prepared daily and sonicated for at least 3 cycles in the provided program to prevent clogging. The control group received DMSO and corn oil, with an injection volume of 5 µL per fish.

DHMEQ was injected daily between 7- and 14-dpci for the 14-dpci time point and between 7- and 30-dpci for the 30-dpci time point isolation. The concentration was determined as 10 mg/kg, with a working solution of 0.8 mg/L and an injection volume of 5-µL per day. A 5 mg stock solution was prepared by dissolving DHMEQ in 1-mL of DMSO: PEG300: Tween80: saline (0.1:0.3:0.05:0.455) and diluted in saline. Control fish received the solvent diluted in saline buffer (0.05 mg/mL) daily.

### Tissue Processing

After isolation, hearts were briefly washed in PBS and fixed overnight in 4% paraformaldehyde at 4°C. The following day, samples were rinsed three times in PBS for 5 minutes each and dehydrated in an ascending ethanol series (70%, 80%, 90%, 96%, 100%) and 100% butanol for 1 hour per step, followed by overnight storage in 100% butanol at 4°C.

Samples were then transferred to paraffin wax (Paraplast Plus, Sigma-Aldrich P-3683) at 65°C. The paraffin was renewed three times at 1-hour intervals, and hearts were embedded in sectioning moulds. After the wax solidified, sections were cut to 7 µm thickness using a Leica microtome and stored in stackable trays.

### Histology

Acid Fuchsin Orange-G (AFOG) staining was performed as previously described by Poss et al. (2002). Hearts were mounted as 1 in 10 sections on Superfrost glass slides and left on a hotplate overnight. Samples were deparaffinised in xylene twice for 5 minutes each, followed by a descending ethanol series (100%, 96%, 90%, 80%, 70%) for 1 minute each. Slides were placed in MilliQ water and refixed in Bouin’s solution for 2 hours at 60°C. After cooling, slides were washed in MilliQ water for 30 minutes, incubated in 1% phosphomolybdic acid for 5 minutes, rinsed with MilliQ water, stained with AFOG solution for 9 minutes, and briefly dehydrated in an ascending ethanol series. AFOG solution was prepared by boiling 1 L of MilliQ water with 5 g of Methyl Blue, followed by the addition of 10 g of Orange-G and 15 g of Acid Fuchsin after cooling. The pH was adjusted to 1.09 using HCl. Following Histoclear treatment for 10 minutes, sections were mounted in DPX mounting medium and left to dry overnight.

### Immunofluorescent Staining

For immunofluorescent staining, at least 3 sections were mounted on slides and left on a hotplate at 37°C overnight. Slides were deparaffinised in two xylene washes (5 minutes each) and rehydrated in a descending ethanol series. Samples were immersed in MilliQ water and pressure-cooked for 8 minutes in Antigen Unmasking Solution (Vector). Slides were cooled and transferred to 0.01% PBS-Tween80 for 5 minutes. Sections were circled with a pap-pen and blocked in TNB buffer for at least 30 minutes at room temperature. Primary antibodies were diluted 1:200 in TNB and applied to sections, which were incubated overnight at room temperature in a humidified box. Slides were washed three times in 0.01% PBS-Tween80 for 5 minutes each, followed by incubation with secondary antibodies (1:250 in TNB) for 2 hours. Sections were counterstained with DAPI, washed three times in 0.01% PBS-Tween80, and mounted in Mowiol for imaging. For wholemount imaging of IgM+ cells, hearts from Tg(*Cau.ighv-ighm:EGFP*)^35^ fish were dissected into ice-cold PBS, fixed in 4% Paraformaldehyde (PFA) at 4°C then further washed in PBS prior to immunostaining. Chicken anti-GFP (1:100, Abcam, ab13970) followed by goat anti-chicken 488 (1:500, Thermo, A11039) antibodies were used following standard procedures.

### RNAscope *In Situ* Hybridization

RNAscope (Advanced Cell Diagnostics, Hayward, CA) was used for *in situ* hybridization on paraffin-embedded sections (7 µm thick). At least 3 sections were mounted per sample and deparaffinised in two xylene washes (5 minutes each), followed by two 2-minute washes in 100% ethanol. After air-drying, sections were treated with hydrogen peroxide for 10 minutes at room temperature and boiled in RNAscope Target Retrieval Solution for 15–18 minutes at 96–102°C. Slides were washed in 100% ethanol and incubated with RNAscope Protease III for 15 minutes at 40°C. Sections were then incubated with the probe mix for 2 hours and treated with amplification buffers and hybridization reagents at 40°C according to the manufacturer’s instructions (RNAscope Multiplex Detection Kit). Hybridization signals were detected using RNAscope Multiplex Fluorescent Detection Reagents v2 with TSA Plus Cyanine 3, Cyanine 5, and Fluorescein fluorophores (Perkin Elmer, NEL744001KT). All *A. mexicanus* custom probes were designed by Advanced Cell Diagnostics (Biotechne). Samples were counterstained with MF-20 (mouse monoclonal antibody for cardiomyocytes) and DAPI, then mounted using gold anti-fade mountant.

### Image Acquisition and Analysis

AFOG images were acquired using a Nikon DS-Fi3 Light Microscope (4x and 10x objectives). All measurements and cell counts were performed blinded. Regeneration capacity was assessed by dividing the total wound area by the total ventricle area, expressed as a percentage or a ratio. Wound closure analysis was performed by measuring the perimeter of open wounds and dividing it by the total ventricle perimeter. The average of three sections with the largest perimeter was used for each heart.

Confocal microscopy was performed using a Zeiss 880. Zebrafish imaging was performed on a Leica TCS SP8 AOBS confocal laser scanning microscope using a 10x/0.4 HC PL APO Dry or 20x/0.75 HC PL APO CS2 Immersion objective. Imaging of AFOG staining on sections was performed on an Evident (Olympus) VS200 slide scanner microscope with colour camera and 20x/0.8 Dry objective.

Fiji/ImageJ software was used for image processing, blinding, wound and ventricular area measurements, and manual quantifications. MF20 was used as a reference to define the wound area. Nuclei colocalised with positive signals were manually quantified using the multipoint tool in Fiji. Counts were normalised to fractional wound area (wound area/total ventricle area). Values from different sections of the same heart were averaged. The number of samples (n) used in each experiment is shown in the figures. Results are expressed as mean ± SD. Statistical analysis of the image quantifications was performed using unpaired t-tests with Welch’s correction and Two-way ANOVA in GraphPad Prism (version 10.1.1 for Windows, Boston, Massachusetts USA, www.graphpad.com).

### Bisque deconvolution and analysis

For the bulk deconvolution, the single-cell FASTQ files were re-mapped to the *Astyanax mexicanus* genome (AstMex3) with the mitogenome (after removing rRNA and tRNA regions) appended. A custom Snakemake pipeline managed the workflow using cell-ranger (v8.0.1) for alignment and quantification. Bulk RNA-seq samples were similarly processed using Fastqc and Trim-galore for quality control and trimming, then mapped with STAR (v2.7.11b) and normalised with DESeq2.

In single cell preprocessing, ambient RNA was removed with decontX and doublets identified via scDblFinder. Quality control thresholds were applied (mitochondrial content <20% for cardiomyocytes and nCount/nFeature within ±2 standard deviations), followed by normalisation using SCTransform (regressing out ribosomal genes) and integration with Harmony. Cell-type annotation was performed using Seurat’s FindAllMarkers, with cxcr4b expression defining broad leukocyte populations and a stricter mitochondrial threshold (<10%) applied to leukocytes; these annotations were then merged into a master reference.

For bulk deconvolution, 100 markers per cell population were identified from the master single-cell dataset using Seurat’s FindAllMarkers. Bisque was then applied—requiring at least 25 unique markers per population—to estimate cell-type proportions in each bulk RNA-seq sample.

Finally, a correlation and pathway analysis were conducted by performing Spearman’s rank correlation between B-cell deconvolution scores and all genes, highlighting those with coefficients >0.7. The ranked gene list was used in FGSEA with WikiPathways to identify the top 20 enriched pathways. Additionally, due to observed differences in B-cell scores at 14 dpci between surface fish and cavefish, these scores were correlated with GSVA summary scores for the “G1 to S Cell Cycle Control” pathway, with Pearson’s correlation coefficient and p-value calculated.

## Data and material availability

Sequencing data are deposited online, all data and material requests should be directed to Mathilda Mommersteeg. The Tg(*Cau.ighv-ighm:EGFP*)^35^ and *rag ^hu^*^1999^ ^36^ mutants are available from Rebecca Richardson.

## Acknowledgements

We thank University of Oxford Biomedical Services and Institute of Developmental and Regenerative Medicine Animal Facility for their continuous efforts and care with animal husbandry. We also thank Carolyn Carr for critical reading of the manuscript and Robin Choudhury for helpful discussions. This work was supported by the Chan Zuckerberg Initiative (MTMM), the European Research Council (ERC) under the European Union’s Horizon 2020 research and innovation program (715895, CAVEHEART, ERC-2016-STG to MTMM), BHF Centre of Regenerative Medicine (RM/13/3/30159), the BHF Centre of Research Excellence Oxford (RE/13/1/30181). ES was funded by an Oxford Medical Research Council Doctoral Training Program and Clarendon Fund (MR/N013468/1), HGP was funded by British Heart Foundation PhD program (FS/17/68/33478), RC was funded by a Novo Nordisk Postdoctoral Fellowship run in partnership with the University of Oxford. JY was funded by University of Oxford Chinese Scholarship Committee. KL was funded by Oxford Medical Research Council Doctoral Training Program (MR/W006731/1). LB, RA and RR were funded by the BBSRC (BB/W001780/1) and acknowledge funding for bioimaging equipment from the MRC (MC_PC_MR/X01391X/1). RR wishes to thank the Wolfson Bioimaging Facility and the Animal Services Unit at the University of Bristol. We also wish to thank Valérie Wittamer, Université Libre de Bruxelles (ULB), for providing IgM and *rag1* lines.

## Author contributions

Conceptualisation: MTMM, ES, HGP. Methodology: MTMM, ES, HGP, WTS, RC, LM, RR. Investigation: ES, HGP, WTS, RC, LB, MN, LOB, RA, JY, KL, ML, RR, MTMM. Formal analysis: ES, HGP, WTS, RC, LB, ML, RR. Writing – Original Draft: ES, MTMM. Writing – Review and Editing: ES, MTMM, HGP, WTS, RC, RR. Supervision: MTMM, RR. Project administration: MTMM. Funding acquisition: MTMM

## Key resources

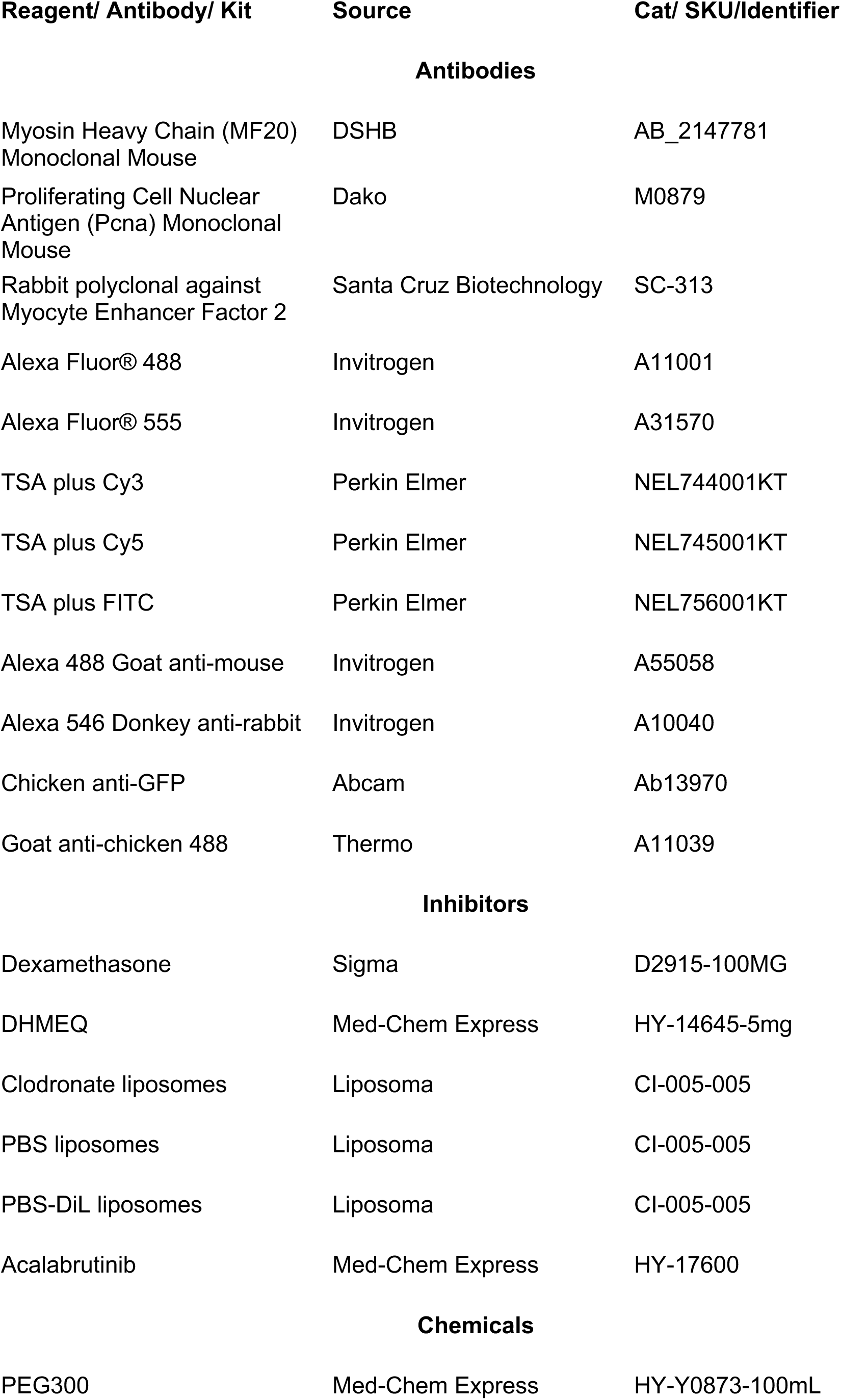
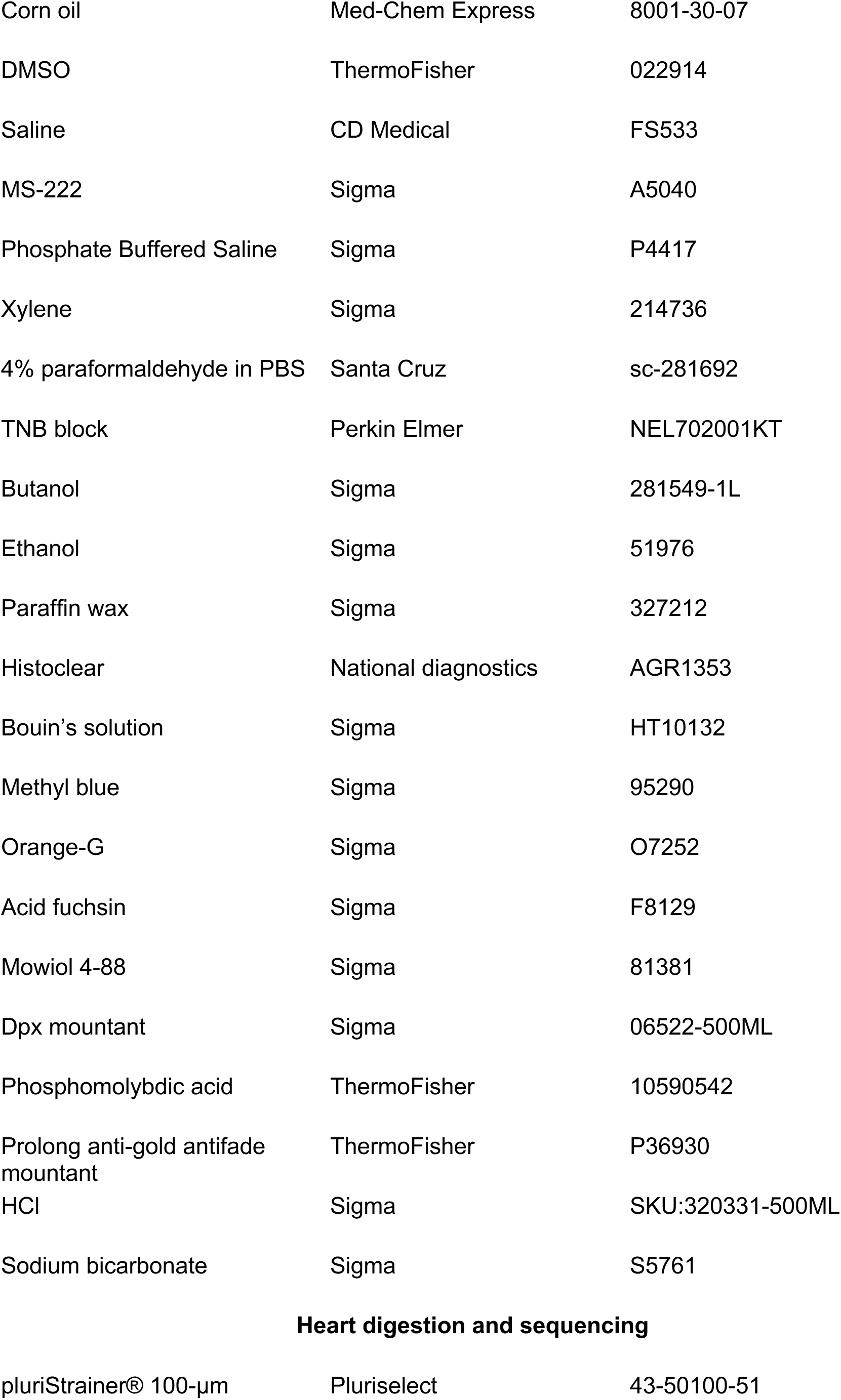
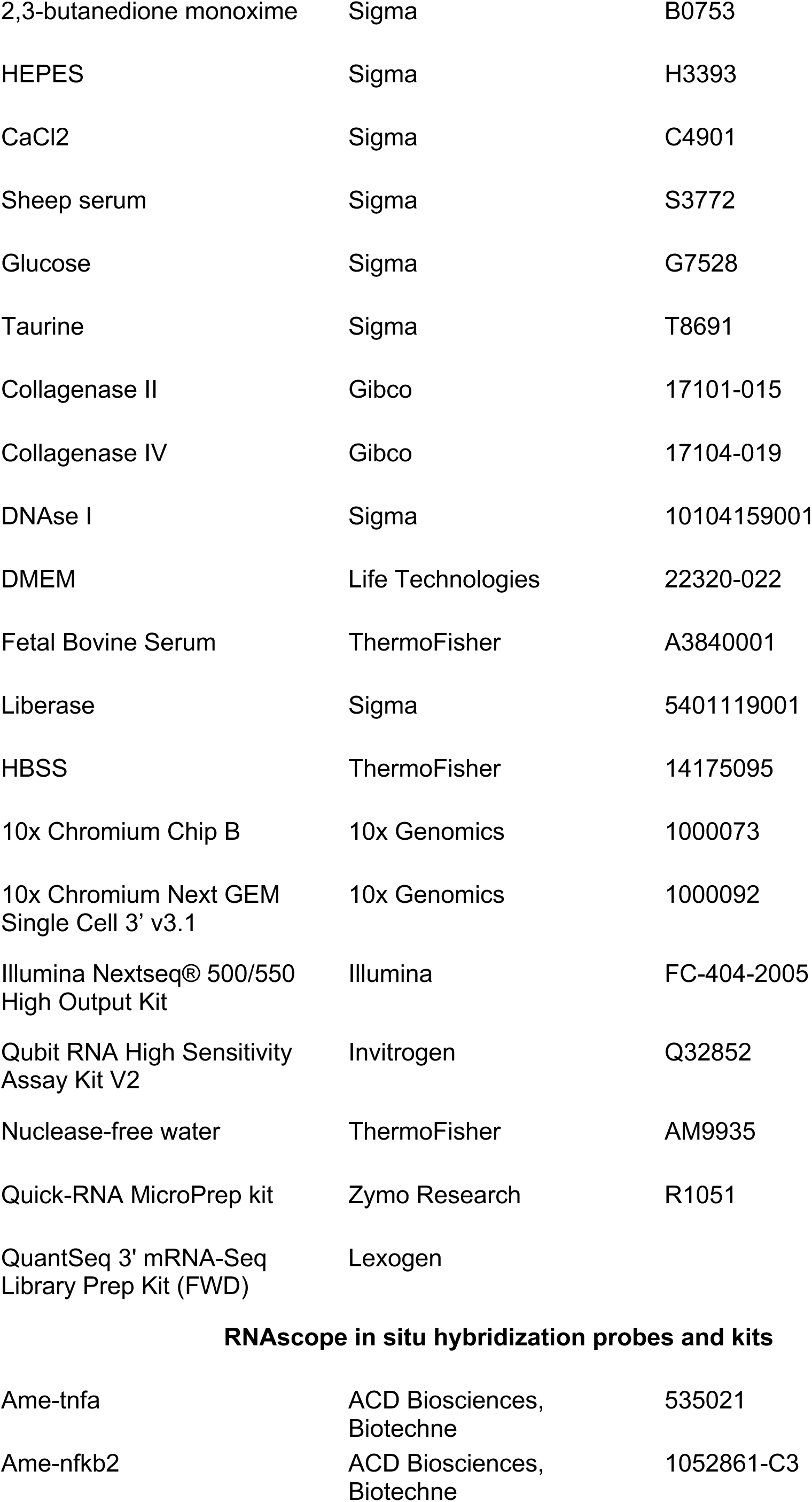
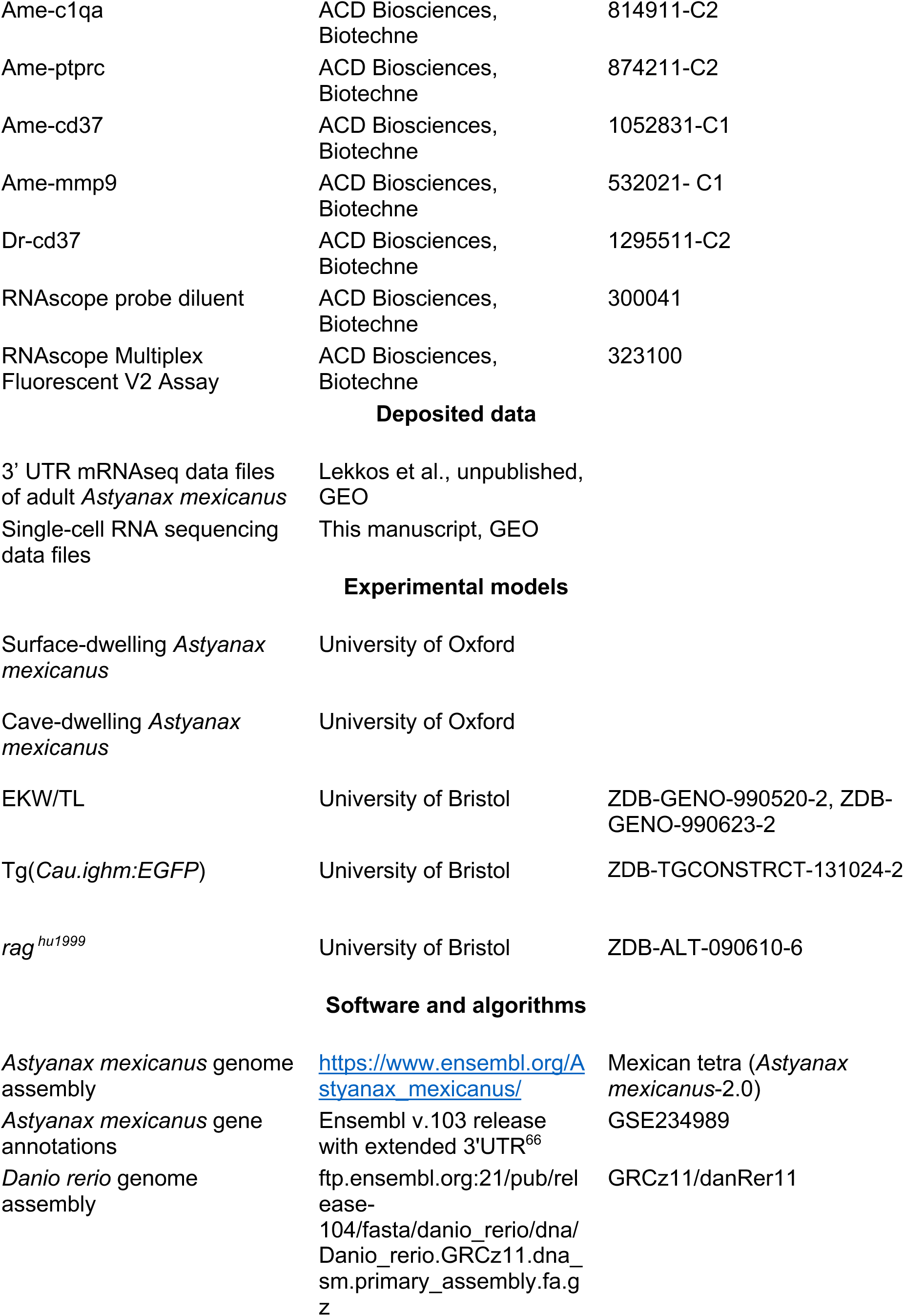
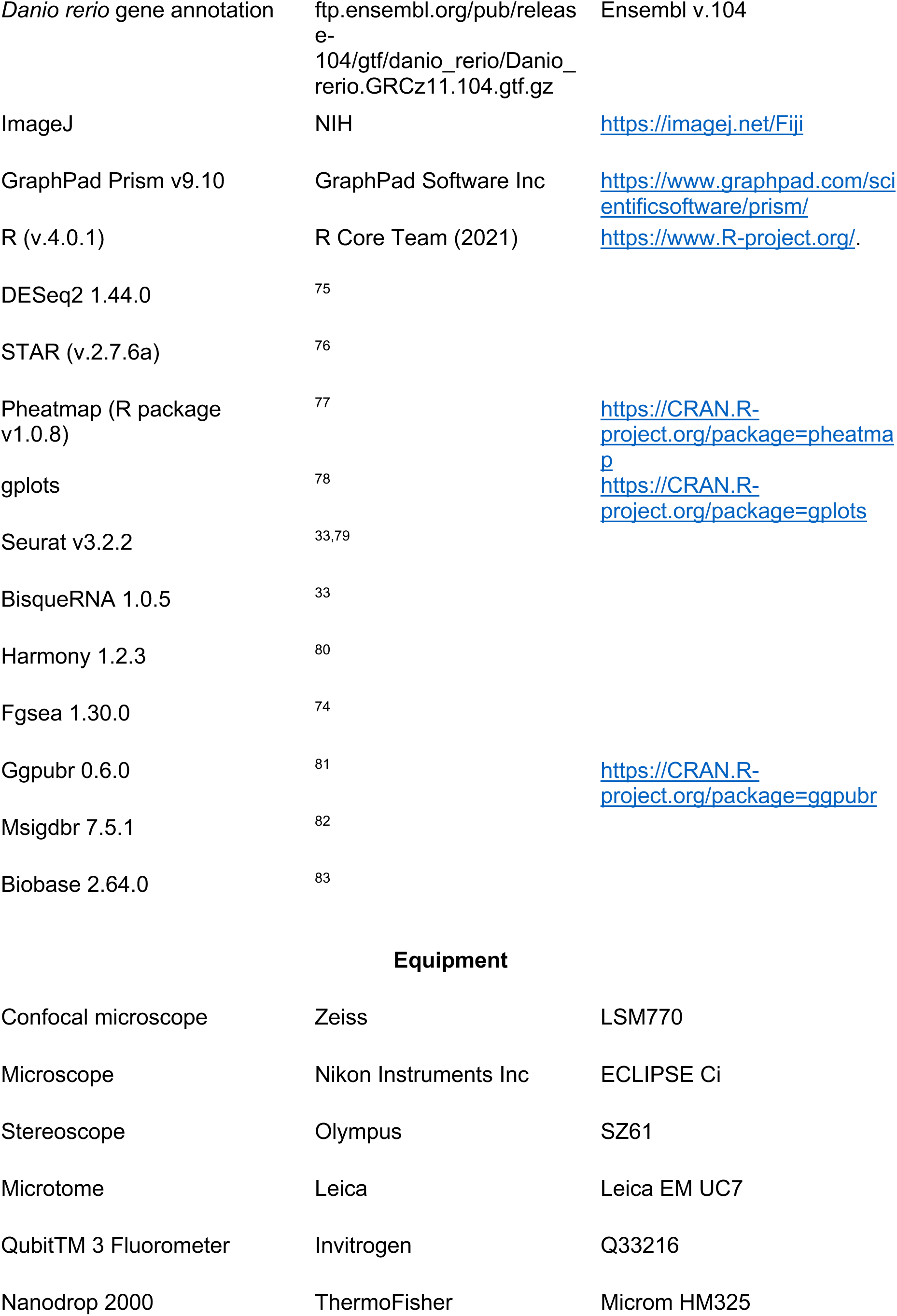
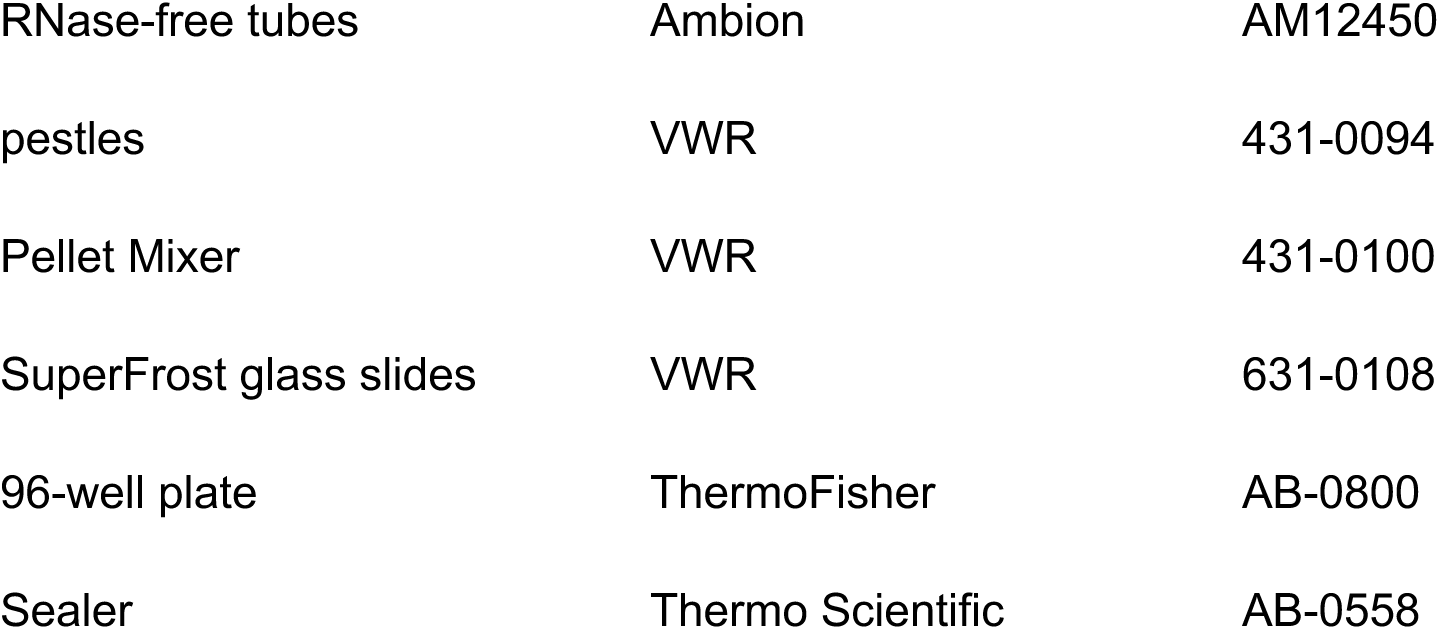

| Reagent/ Antibody/ Kit | Source | Cat/ SKU/Identifier |
| --- | --- | --- |
| **Antibodies** |  |  |
| Myosin Heavy Chain (MF20) Monoclonal Mouse | DSHB | AB_2147781 |
| Proliferating Cell Nuclear Antigen (Pcna) Monoclonal Mouse | Dako | M0879 |
| Rabbit polyclonal against Myocyte Enhancer Factor 2 | Santa Cruz Biotechnology | SC-313 |
| Alexa Fluor® 488 | Invitrogen | A11001 |
| Alexa Fluor® 555 | Invitrogen | A31570 |
| TSA plus Cy3 | Perkin Elmer | NEL744001KT |
| TSA plus Cy5 | Perkin Elmer | NEL745001KT |
| TSA plus FITC | Perkin Elmer | NEL756001KT |
| Alexa 488 Goat anti-mouse | Invitrogen | A55058 |
| Alexa 546 Donkey anti-rabbit | Invitrogen | A10040 |
| Chicken anti-GFP | Abcam | Ab13970 |
| Goat anti-chicken 488 | Thermo | A11039 |
| **Inhibitors** |  |  |
| Dexamethasone | Sigma | D2915-100MG |
| DHMEQ | Med-Chem Express | HY-14645-5mg |
| Clodronate liposomes | Liposoma | CI-005-005 |
| PBS liposomes | Liposoma | CI-005-005 |
| PBS-DiL liposomes | Liposoma | CI-005-005 |
| Acalabrutinib | Med-Chem Express | HY-17600 |
| **Chemicals** |  |  |
| PEG300 | Med-Chem Express | HY-Y0873-100mL |
| Corn oil | Med-Chem Express | 8001-30-07 |
| DMSO | ThermoFisher | 022914 |
| Saline | CD Medical | FS533 |
| MS-222 | Sigma | A5040 |
| Phosphate Buffered Saline | Sigma | P4417 |
| Xylene | Sigma | 214736 |
| 4% paraformaldehyde in PBS | Santa Cruz | sc-281692 |
| TNB block | Perkin Elmer | NEL702001KT |
| Butanol | Sigma | 281549-1L |
| Ethanol | Sigma | 51976 |
| Paraffin wax | Sigma | 327212 |
| Histoclear | National diagnostics | AGR1353 |
| Bouin's solution | Sigma | HT10132 |
| Methyl blue | Sigma | 95290 |
| Orange-G | Sigma | O7252 |
| Acid fuchsin | Sigma | F8129 |
| Mowiol 4-88 | Sigma | 81381 |
| Dpx mountant | Sigma | 06522-500ML |
| Phosphomolybdic acid | ThermoFisher | 10590542 |
| Prolong anti-gold antifade mountant | ThermoFisher | P36930 |
| HCl | Sigma | SKU:320331-500ML |
| Sodium bicarbonate | Sigma | S5761 |
| **Heart digestion and sequencing** |  |  |
| pluriStrainer® 100-µm | Pluriselect | 43-50100-51 |
| 2,3-butanedione monoxime | Sigma | B0753 |
| HEPES | Sigma | H3393 |
| CaCl<sub>2</sub> | Sigma | C4901 |
| Sheep serum | Sigma | S3772 |
| Glucose | Sigma | G7528 |
| Taurine | Sigma | T8691 |
| Collagenase II | Gibco | 17101-015 |
| Collagenase IV | Gibco | 17104-019 |
| DNAse I | Sigma | 10104159001 |
| DMEM | Life Technologies | 22320-022 |
| Fetal Bovine Serum | ThermoFisher | A3840001 |
| Liberase | Sigma | 5401119001 |
| HBSS | ThermoFisher | 14175095 |
| 10x Chromium Chip B | 10x Genomics | 1000073 |
| 10x Chromium Next GEM Single Cell 3' v3.1 | 10x Genomics | 1000092 |
| Illumina Nextseq® 500/550 High Output Kit | Illumina | FC-404-2005 |
| Qubit RNA High Sensitivity Assay Kit V2 | Invitrogen | Q32852 |
| Nuclease-free water | ThermoFisher | AM9935 |
| Quick-RNA MicroPrep kit | Zymo Research | R1051 |
| QuantSeq 3' mRNA-Seq Library Prep Kit (FWD) | Lexogen |  |
| **RNAscope in situ hybridization probes and kits** |  |  |
| Ame-tnfa | ACD Biosciences, Biotechne | 535021 |
| Ame-nfkb2 | ACD Biosciences, Biotechne | 1052861-C3 |
| Ame-c1qa | ACD Biosciences,<br>Biotechnne | 814911-C2 |
| Ame-ptprc | ACD Biosciences,<br>Biotechnne | 874211-C2 |
| Ame-cd37 | ACD Biosciences,<br>Biotechnne | 1052831-C1 |
| Ame-mmp9 | ACD Biosciences,<br>Biotechnne | 532021- C1 |
| Dr-cd37 | ACD Biosciences,<br>Biotechnne | 1295511-C2 |
| RNAscope probe diluent | ACD Biosciences,<br>Biotechnne | 300041 |
| RNAscope Multiplex<br>Fluorescent V2 Assay | ACD Biosciences,<br>Biotechnne | 323100 |

Deposited data
|  |  |
| --- | --- |
| 3' UTR mRNAseq data files<br>of adult *Astyanax mexicanus* | Lekkos et al., unpublished,<br>GEO |
| Single-cell RNA sequencing<br>data files | This manuscript, GEO |
| Experimental models |  |  |
| Surface-dwelling *Astyanax*<br>*mexicanus* | University of Oxford |  |
| Cave-dwelling *Astyanax*<br>*mexicanus* | University of Oxford |  |
| EKW/TL | University of Bristol | ZDB-GENO-990520-2, ZDB-GENO-990623-2 |
| Tg(*Cau.ighm:EGFP*) | University of Bristol | ZDB-TGCONSTRCT-131024-2 |
| *rag*<sup>hu1999</sup> | University of Bristol | ZDB-ALT-090610-6 |
| Software and algorithms |  |  |
| *Astyanax mexicanus* genome<br>assembly | https://www.ensembl.org/Astyanax_mexicanus/ | Mexican tetra (*Astyanax*<br>*mexicanus*-2.0) |
| *Astyanax mexicanus* gene<br>annotations | Ensembl v.103 release<br>with extended 3'UTR<sup>66</sup> | GSE234989 |
| *Danio rerio* genome<br>assembly | ftp.ensembl.org:21/pub/rel<br>ease-104/fasta/danio_rerio/dna/<br>Danio_rerio.GRCz11.dna_<br>sm.primary_assembly.fa.g<br>z | GRCz11/danRer11 |
| *Danio rerio* gene annotation | ftp.ensembl.org/pub/release-104/gtf/danio_rerio/Danio_rerio.GRCz11.104.gtf.gz | Ensembl v.104 |
| ImageJ | NIH | https://imagej.net/Fiji |
| GraphPad Prism v9.10 | GraphPad Software Inc | https://www.graphpad.com/scientificsoftware/prism/ |
| R (v.4.0.1) | R Core Team (2021) | https://www.R-project.org/. |
| DESeq2 1.44.0 | 75 |  |
| STAR (v.2.7.6a) | 76 |  |
| Pheatmap (R package v1.0.8) | 77 | https://CRAN.R-project.org/package=pheatmap |
| gplots | 78 | https://CRAN.R-project.org/package=gplots |
| Seurat v3.2.2 | 33,79 |  |
| BisqueRNA 1.0.5 | 33 |  |
| Harmony 1.2.3 | 80 |  |
| Fgsea 1.30.0 | 74 |  |
| Ggpubr 0.6.0 | 81 | https://CRAN.R-project.org/package=ggpubr |
| Msigdb 7.5.1 | 82 |  |
| Biobase 2.64.0 | 83 |  |
| Equipment |  |  |
| Confocal microscope | Zeiss | LSM770 |
| Microscope | Nikon Instruments Inc | ECLIPSE Ci |
| Stereoscope | Olympus | SZ61 |
| Microtome | Leica | Leica EM UC7 |
| Qubit™ 3 Fluorometer | Invitrogen | Q33216 |
| Nanodrop 2000 | ThermoFisher | Microm HM325 |
| RNase-free tubes | Ambion | AM12450 |
| pestles | VWR | 431-0094 |
| Pellet Mixer | VWR | 431-0100 |
| SuperFrost glass slides | VWR | 631-0108 |
| 96-well plate | ThermoFisher | AB-0800 |
| Sealer | Thermo Scientific | AB-0558 |

## Supplementary Information

**Supplementary Fig. 1.**
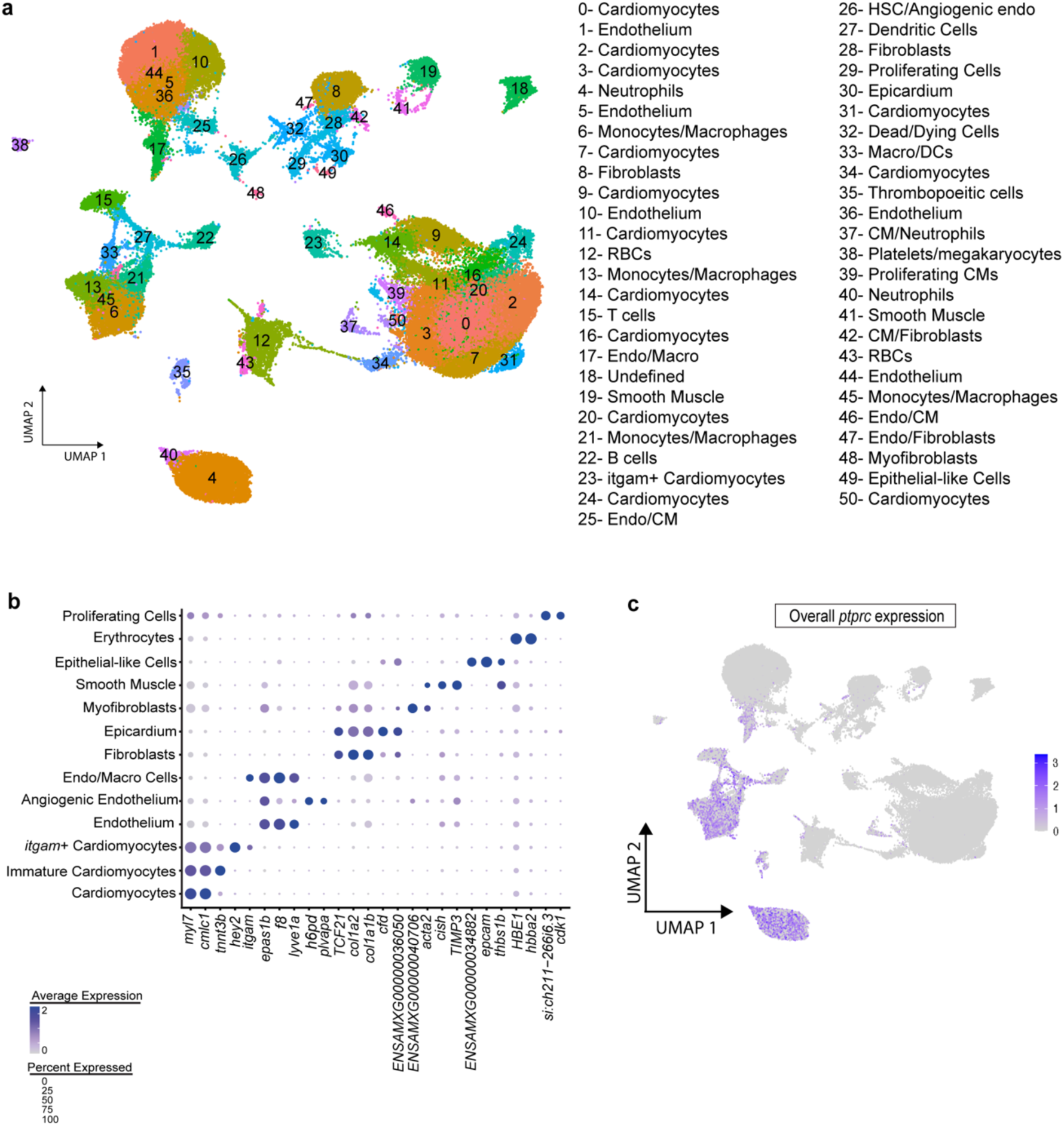
Single cell profiling of *Astyanax mexicanus* heart cell populations. a UMAP visualisation of all annotated cell populations in the *A. mexicanus* heart. b Dot plot displaying the expression of defining marker genes for each annotated cell type. c UMAP plot highlighting the expression of *ptprc*, across all cell populations in the heart.

**Supplementary Fig. 2.**
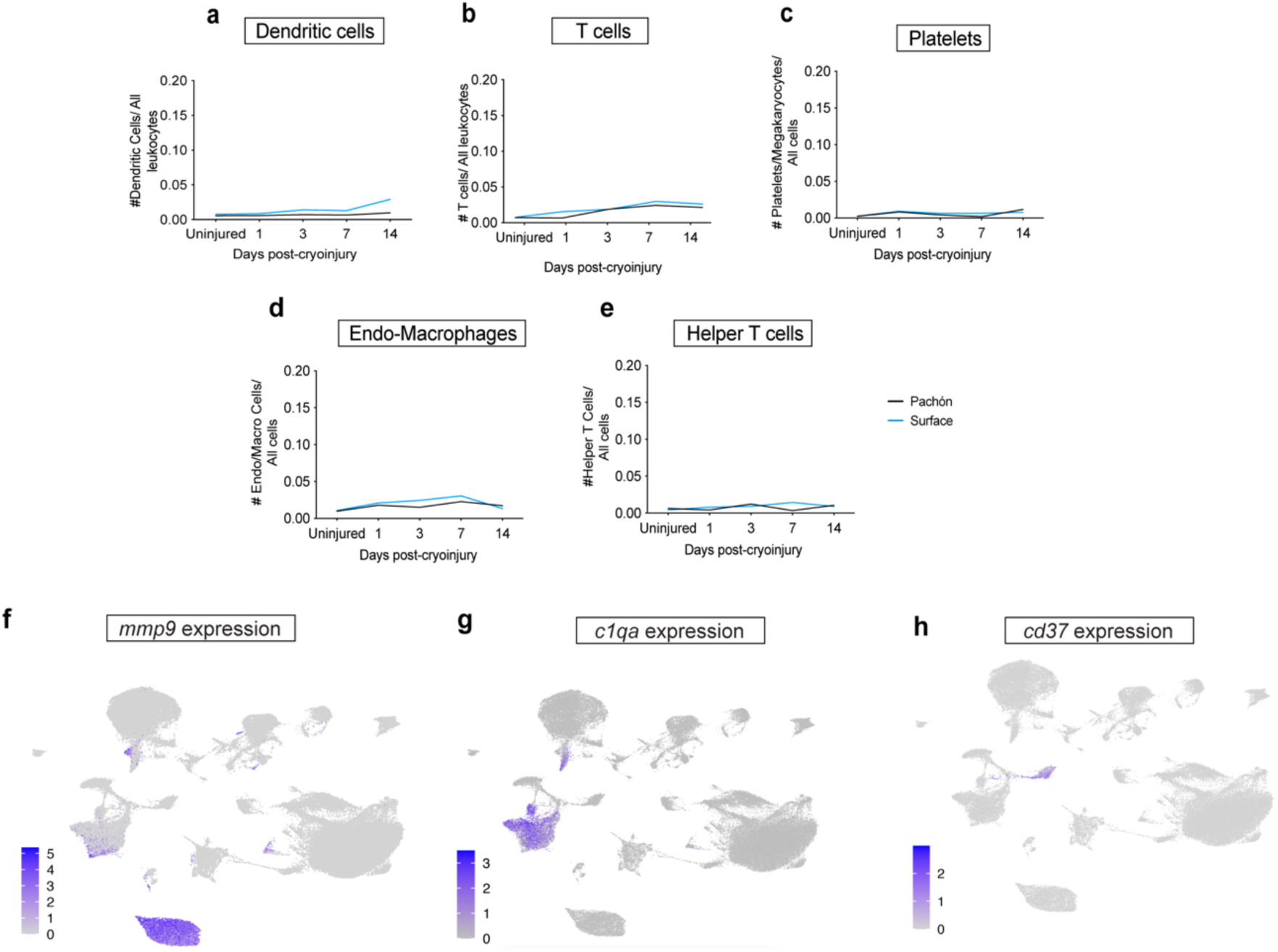
Temporal dynamics of immune cell populations in *A. mexicanus* heart regeneration. **a-c** Differential proportion analysis of dendritic cells, (**a**), T cells (**b**), platelets (**c**), endo-macrophage cells (**d**), and helper T cells (**e**) at different time points in Pachón and surface fish hearts, revealing changes in cell abundance during regeneration. **f-h** UMAPs representing the overall expression of *mm p 9*, *c1qa*, and *cd37* marker genes, respectively.

**Supplementary Fig. 3.**
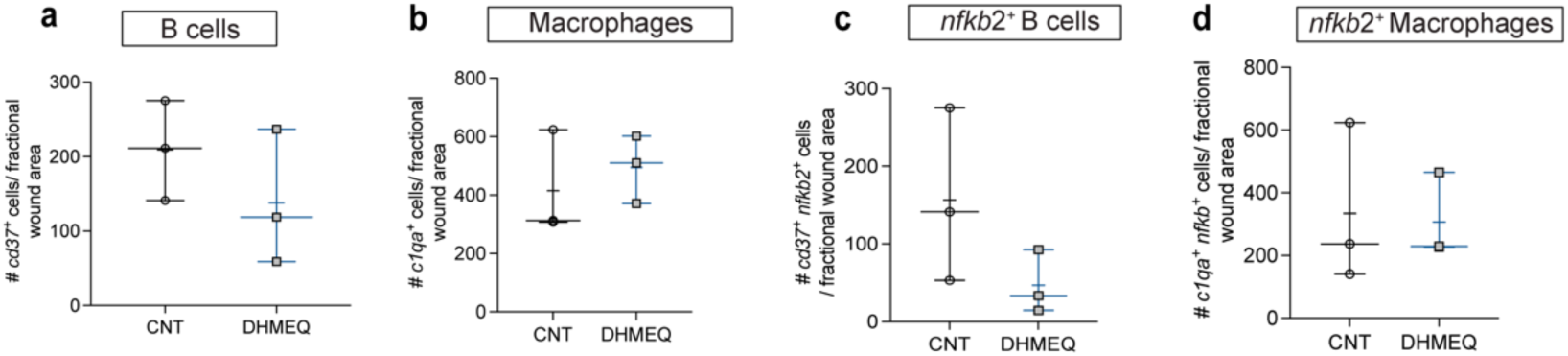
Immune cell responses in wound healing after cardiac injury. a-d Quantification of *cd37*^+^ (a), *c1qa^+^* (b), *cd37*^+^*nfkb2*^+^ (c), and *c1qa*^+^*nfkb2*^+^ (d) cells in the wound at 14-dpci in control and DHMEQ-treated surface fish hearts, showing no significant differences between groups.

**Supplementary Fig. 4.**
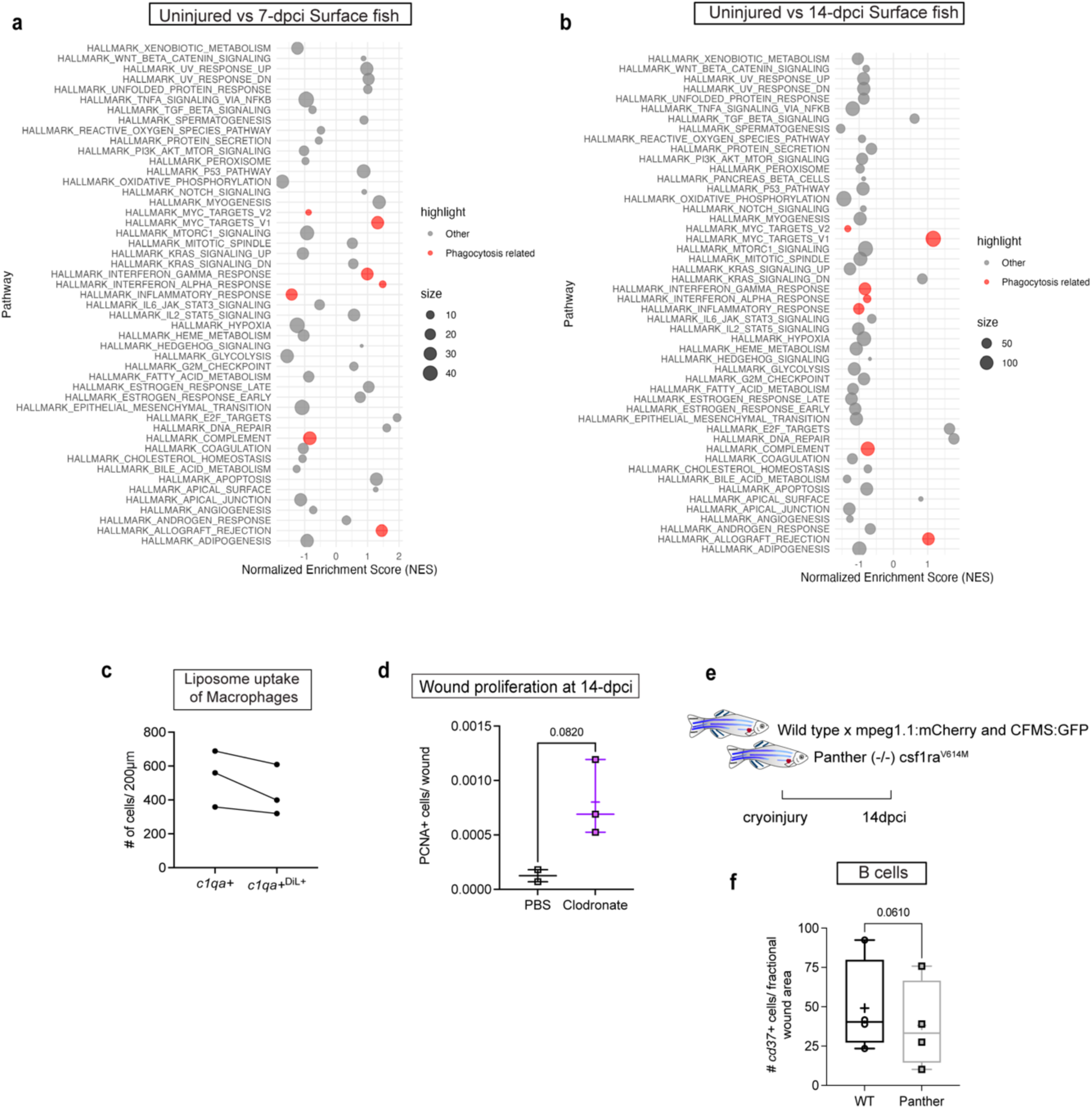
Macrophage-mediated pathways and cellular responses during heart regeneration. **a-b** Gene Set Enrichment Analysis (GSEA) results showing Hallmark pathways that reached the FDR significance threshold (FDR=0.25) for 7-dpci (**a**) and 14-dpci (**b**) surface fish macrophages compared to uninjured controls. Phagocytosis-related pathways are highlighted in red, while other pathways are shown in grey. **c** Co-localisation analysis of total *c1qa*^+^ counts with *c1qa*^+^/DiL^+^ counts (n=3). **d** Quantification of pcna-positive proliferating cells in the wound, showing a significant difference between PBS Control and Clodronate-treated groups, indicating that macrophage depletion inhibits cell influx to the wound (unpaired t-test). **e** RNAscope analysis of cryoinjured hearts from Wild Type and Panther (*csf1ra*^-/-^) zebrafish at 14-dpci. f Quantification of *cd37*^+^ cells showing no difference in B cell response between the control and Panther zebrafish.

**Supplementary Fig. 5.**
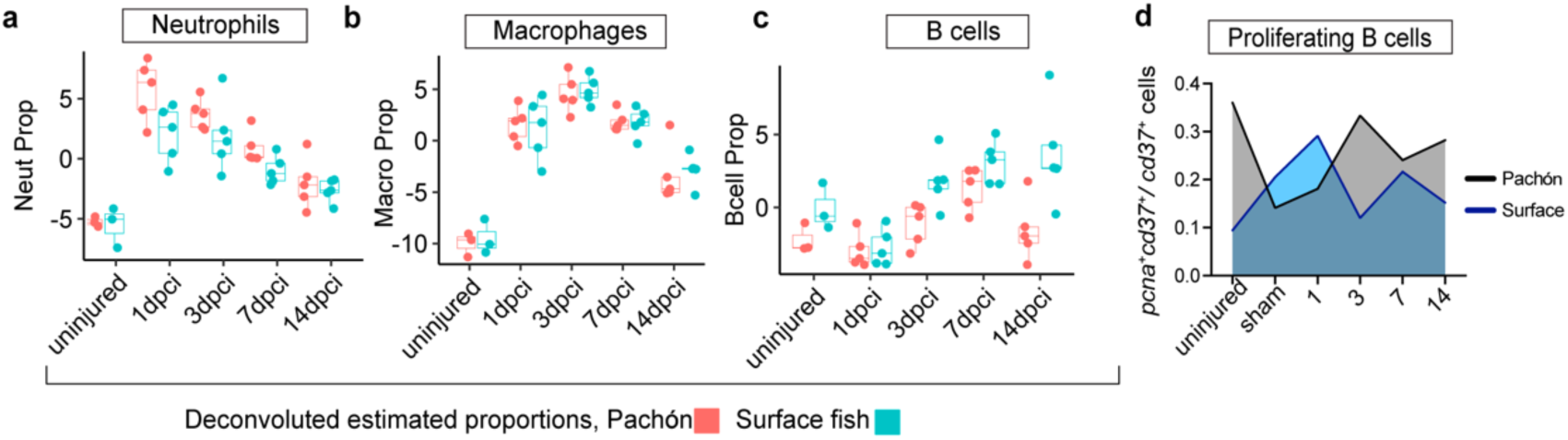
Temporal dynamics of immune cell populations during heart regeneration. **a-d** Deconvoluted differential proportion analysis showing the relative abundance of neutrophils (**a**), macrophages (**b**), and B cells (**c**) at 1, 3, 7, 14-dpci and uninjured time points in both surface and Pachón fish morphs. d Proportion of proliferating B cells (*pcna*^+^*cd37*^+^) to total B cells (*cd37*^+^) in surface fish and Pachón ventricles, derived from scRNA-seq data.

**Supplementary Fig. 6.**
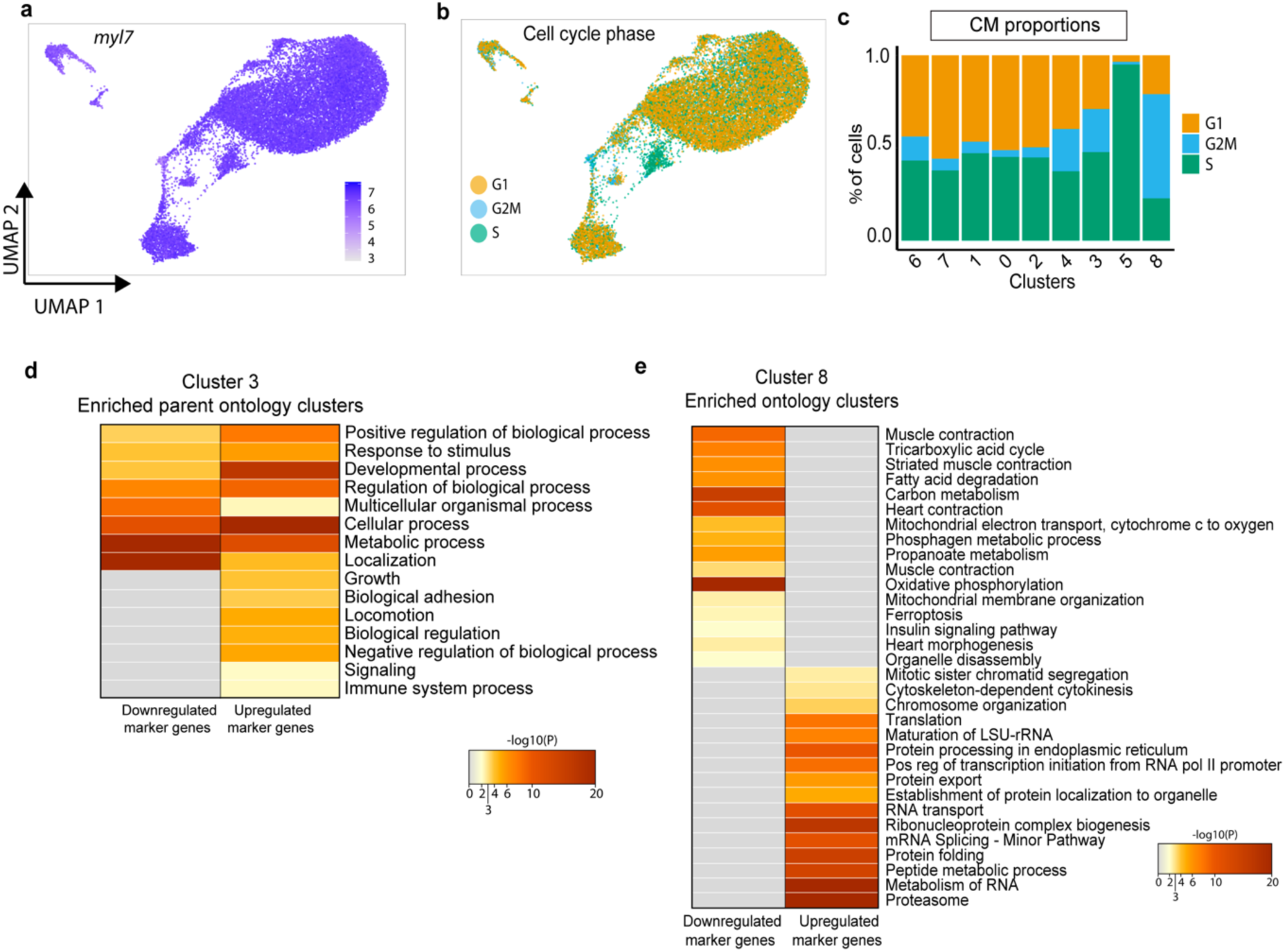
Cardiomyocyte heterogeneity and cell cycle dynamics during heart regeneration. **a** UMAP of re-clustered cardiomyocyte scRNA-seq data, showing widespread expression of *myl7*. **b** Border zone cardiomyocytes exhibit high expression of G2M, and S cell cycle phases compared to other sub-cardiomyocyte populations. **c** Differential proportion analysis of cardiomyocytes, showing the percentage of cells in G1, G2M, and S phases across different cardiomyocyte clusters. **d** GO terms for cluster 3, showing strong upregulation in regulation and signaling, growth and development, and structural and adhesion processes. **e** GO terms for cluster 8, showing strong upregulation in chromosome and cell division processes, RNA processing, and protein and peptide metabolism.

---

**Supplementary table.** Overview of statistical tests and values displayed on the figures.

| Figure | Analysis | Comparisons | Time point | p value | Test type/ comparisons |
| --- | --- | --- | --- | --- | --- |
| Fig. 1d | DPA overall leukocytes | Pachón - surface | 7 | 0.0199 | Permutation analysis |
| Fig. 1f | #ptprc cells | Pachón - surface | 14 | 0.0037 | Šídák's multiple comparisons test |
| Fig. 1f | #ptprc cells | Pachón | Uninjured vs. 3 | 0.0001 | Tukey's multiple comparisons test |
| Fig. 1f | #ptprc cells | Pachón | 3 vs. 7 | 0.0445 | Tukey's multiple comparisons test |
| Fig. 1f | #ptprc cells | Pachón | 3 vs. 14 | 0.0004 | Tukey's multiple comparisons test |
| Fig. 1f | #ptprc cells | Pachón | 3 vs. 30 | 0.0036 | Tukey's multiple comparisons test |
| Fig. 1f | #ptprc cells | Pachón | 3 vs. 60 | 0.0002 | Tukey's multiple comparisons test |
| Fig. 1f | #ptprc cells | Surface fish | Uninjured vs. 1 | <0.0001 | Tukey's multiple comparisons test |
| Fig. 1f | #ptprc cells | Surface fish | Uninjured vs. 3 | 0.0003 | Tukey's multiple comparisons test |
| Fig. 1f | #ptprc cells | Surface fish | Uninjured vs. 7 | 0.0229 | Tukey's multiple comparisons test |
| Fig. 1f | #ptprc cells | Surface fish | Uninjured vs. 14 | 0.0027 | Tukey's multiple comparisons test |
| Fig. 1f | #ptprc cells | Surface fish | Uninjured vs. 30 | 0.0227 | Tukey's multiple comparisons test |
| Fig. 1f | #ptprc cells | Surface fish | 1 vs. 30 | 0.0331 | Tukey's multiple comparisons test |
| Fig. 1f | #ptprc cells | Surface fish | 1 vs. 60 | 0.0052 | Tukey's multiple comparisons test |
| Fig. 1i | scarring upon Dexamethasone | Dexamethasone vs water control | 14 | 0.0491 | Unpaired t test, Two-tailed |
| Fig. 2c | DPA- neutrophils | Pachón - surface | 3 | 0.0007 | Permutation analysis |
| Fig. 2d | DPA- monocytes & macrophages | Pachón - surface | 7 | 0.0109 | Permutation analysis |
| Fig. 2e | DPA B cells | Pachón - surface | 14 | 0.0143 | Permutation analysis |
| Fig. 2g | #mmp9 cells | Pachón | uninjured<br>vs. 1 | 0.00<br>21 | Tukey's multiple<br>comparisons test |
| Fig. 2g | #mmp9 cells | Pachón | 1 vs. 7 | 0.00<br>17 | Tukey's multiple<br>comparisons test |
| Fig. 2g | #mmp9 cells | Pachón | 1 vs. 14 | 0.01<br>04 | Tukey's multiple<br>comparisons test |
| Fig. 2g | #mmp9 cells | Pachón | 1 vs. 30 | 0.00<br>07 | Tukey's multiple<br>comparisons test |
| Fig. 2g | #mmp9 cells | Pachón | 1 vs. 60 | 0.00<br>22 | Tukey's multiple<br>comparisons test |
| Fig. 2i | #c1qa cells | Pachón - surface | 14 | 0.02<br>61 | F-test to compare<br>variances |
| Fig. 2i | #c1qa cells | Pachón | uninjured<br>vs. 3 | 0.04 | Tukey's multiple<br>comparisons test |
| Fig. 2i | #c1qa cells | Pachón | 3 vs. 30 | 0.02<br>81 | Tukey's multiple<br>comparisons test |
| Fig. 2i | #c1qa cells | Pachón | 7 vs. 30 | 0.04<br>61 | Tukey's multiple<br>comparisons test |
| Fig. 2i | #c1qa cells | Surface | uninjured<br>vs. 3 | 0.03<br>46 | Tukey's multiple<br>comparisons test |
| Fig. 2i | #c1qa cells | Surface | uninjured<br>vs. 7 | 0.00<br>33 | Tukey's multiple<br>comparisons test |
| Fig. 2i | #c1qa cells | Surface | uninjured<br>vs. 14 | 0.04<br>32 | Tukey's multiple<br>comparisons test |
| Fig. 2i | #c1qa cells | Surface | 7 vs. 30 | 0.02<br>42 | Tukey's multiple<br>comparisons test |
| Fig. 2i | #c1qa cells | Surface | 7 vs. 60 | 0.03<br>77 | Tukey's multiple<br>comparisons test |
| Fig. 2k | #cd37 cells | Pachón - surface | 30 | <0.0<br>001 | Šidák's multiple<br>comparisons test |
| Fig. 2k | #cd37 cells | Surface | uninjured<br>vs. 30 | <0.0<br>001 | Tukey's multiple<br>comparisons test |
| Fig. 2k | #cd37 cells | Surface | 7 vs. 30 | <0.0<br>001 | Tukey's multiple<br>comparisons test |
| Fig. 2k | #cd37 cells | Surface | 14 vs. 30 | 0.00<br>16 | Tukey's multiple<br>comparisons test |
| Fig. 2k | #cd37 cells | Surface | 30 vs. 60 | <0.0<br>001 | Tukey's multiple<br>comparisons test |
| Fig. 3k | scarring upon DHMEQ | Surface control vs<br>DHMEQ | 30 | 0.33<br>54 | Unpaired t-test, two-tailed |
| Fig. 4c | #c1qa upon<br>Dexamethasone | Surface control vs<br>Dexamethasone | 14 | 0.59<br>71 | Unpaired t-test, two-tailed |
| Fig. 4f | #c1qa upon Clodronate | Surface control vs<br>Clodronate | 14 | 0.00<br>84 | Unpaired t-test, two-tailed |
| Fig. 4h | scarring upon Clodronate | Surface control vs Clodronate | 30 | 0.0089 | Unpaired t-test, two-tailed |
| Fig. 4j | #cd37 upon Clodronate | Surface control vs Clodronate | 14 | 0.6911 | Unpaired t-test, two-tailed |
| Fig. 5a | #cd37 upon Dexamethasone | Surface control vs Dexamethasone | 14 | 0.018 | Unpaired t-test, two-tailed |
| Fig. 5b | #cd37 upon Dexamethasone | Surface control vs Dexamethasone | 30 | 0.0051 | Unpaired t-test, two-tailed |
| Fig. 5d | #cd37 upon Acalabrutinib | Surface control vs Dexamethasone | 14 | 0.0005 | Unpaired t-test, two-tailed |
| Fig. 5f | scarring upon Acalabrutinib | Surface control vs Acalabrutinib | 30 | 0.0233 | Unpaired t-test, two-tailed |
| Fig. 6f | BZ CM proliferation rate upon Dexamethasone | Surface control vs Dexamethasone | 14 | 0.0055 | Unpaired t-test, two-tailed |
| Fig. 6g | BZ CM proliferation rate upon Clodronate | Surface control vs Clodronate | 14 | 0.6745 | Unpaired t-test, two-tailed |
| Fig. 6h | BZ CM proliferation rate upon Acalabrutinib | Surface control vs Acalabrutinib | 14 | 0.0506 | Unpaired t-test, two-tailed |
| Fig. 6j | BZ CM proliferation rate upon Acalabrutinib | Surface control vs Acalabrutinib | 5 | 0.8629 | Unpaired t-test, two-tailed |
| Fig.7c | #IgM cells | Tg(*Cau.ighv-ighm:EGFP*) | uninjured vs 14 | 0.1198 | One-way ANOVA |
| Fig.7c | #IgM cells | Tg(*Cau.ighv-ighm:EGFP*) | uninjured vs 21 | 0.0265 | One-way ANOVA |
| Fig.7f | scarring | WT vs RAG | 21 | 0.0289 | Unpaired t-test, two-tailed |
| Fig.7f | scarring | WT vs RAG | 60 | 0.0007 | Unpaired t-test, two-tailed |
| Fig.7g | BZ CM proliferation rate upon Acalabrutinib | WT vs RAG | 14 | 0.0261 | Unpaired t-test, two-tailed |
| Supplementary Fig. 3a | #cd37 upon DHMEQ | Surface control vs DHMEQ | 14 | 0.0508 | Unpaired t-test, two-tailed |
| Supplementary Fig. 3b | #c1qa upon DHMEQ | Surface control vs DHMEQ | 14 | 0.5681 | Unpaired t-test, two-tailed |
| Supplementary Fig. 3c | #nfkb2+cd37 upon DHMEQ | Surface control vs DHMEQ | 14 | 0.3326 | Unpaired t-test, two-tailed |
| Supplementary Fig. 3d | #nfkb2+c1qa upon DHMEQ | Surface control vs DHMEQ | 14 | 0.9092 | Unpaired t-test, two-tailed |
| Supplementary Fig. 4d | Overall wound proliferation upon Clodronate | Surface control vs Clodronate | 14 | 0.082 | Unpaired t-test, two-tailed |
| Supplementary Fig. 4f | #cd37 cells | WT vs Panther | 14 | 0.061 | Unpaired t-test, two-tailed |

## References

1. Poss, K. D., Wilson, L. G. & Keating, M. T. Heart Regeneration in Zebrafish. Science (1979) 298, 2188–2190 (2002).

2. Witman, N., Murtuza, B., Davis, B., Arner, A. & Morrison, J. I. Recapitulation of developmental cardiogenesis governs the morphological and functional regeneration of adult newt hearts following injury. Dev Biol 354, 67–76 (2011).

3. Chablais, F., Veit, J., Rainer, G. & Jawiska, A. The zebrafish heart regenerates after cryoinjury-induced myocardial infarction. BMC Dev Biol 11, (2011).

4. Lafontant, P. J. et al. The Giant Danio (D. Aequipinnatus) as A Model of Cardiac Remodeling and Regeneration. Anatomical Record 295, 234–248 (2012).

5. Simões, F. C. & Riley, P. R. Immune cells in cardiac repair and regeneration. Development 149, (2022).

6. Adrover, J. M. et al. A Neutrophil Timer Coordinates Immune Defense and Vascular Protection. Immunity 50, 390–402.e10 (2019).

7. Swirski, F. K. & Nahrendorf, M. Cardioimmunology: the immune system in cardiac homeostasis and disease. Nature Reviews Immunology vol. 18 733– 744 Preprint at 10.1038/s41577-018-0065-8 (2018).

8. Porrello, E. R. et al. Transient regenerative potential of the neonatal mouse heart. Science (1979) 331, 1078–1080 (2011).

9. Aurora, A. B. et al. Macrophages are required for neonatal heart regeneration. Journal of Clinical Investigation 124, 1382–1392 (2014).

10. Foglia, M. J. & Poss, K. D. Building and re-building the heart by cardiomyocyte proliferation. Development (Cambridge) vol. 143 729–740 Preprint at 10.1242/dev.132910 (2016).

11. Godwin, J. W., Pinto, A. R. & Rosenthal, N. A. Macrophages are required for adult salamander limb regeneration. Proc Natl Acad Sci U S A 110, 9415– 9420 (2013).

12. Sanz-Morejón, A. et al. Wilms Tumor 1b Expression Defines a Pro-regenerative Macrophage Subtype and Is Required for Organ Regeneration in the Zebrafish. Cell Rep 28, 1296–1306.e6 (2019).

13. Lavine, K. J. et al. Distinct macrophage lineages contribute to disparate patterns of cardiac recovery and remodeling in the neonatal and adult heart. Proc Natl Acad Sci U S A 111, 16029–16034 (2014).

14. Wynn, T. A. Cellular and molecular mechanisms of fibrosis. Journal of Pathology vol. 214 199–210 Preprint at 10.1002/path.2277 (2008).

15. Bermea, K., Bhalodia, A., Huff, A., Rousseau, S. & Adamo, L. The Role of B Cells in Cardiomyopathy and Heart Failure. Current Cardiology Reports vol. 24 935–946 Preprint at 10.1007/s11886-022-01722-4 (2022).

16. Goodchild, T. T. et al. Bone Marrow-Derived B Cells Preserve Ventricular Function After Acute Myocardial Infarction. JACC Cardiovasc Interv 2, 1005– 1016 (2009).

17. Tan, Y., Duan, X., Wang, B., Liu, X. & Zhan, Z. Murine neonatal cardiac B cells promote cardiomyocyte proliferation and heart regeneration. NPJ Regen Med 8, (2023).

18. Wu, L. et al. Recent Advances on Phagocytic B Cells in Teleost Fish. Frontiers in Immunology vol. 11 Preprint at 10.3389/fimmu.2020.00824 (2020).

19. Stockdale, W. T. et al. Heart Regeneration in the Mexican Cavefish. Cell Rep 25, 1997–2007.e7 (2018).

20. Peuß, R. et al. Adaptation to low parasite abundance affects immune investment and immunopathological responses of cavefish. Nat Ecol Evol 4, 1416–1430 (2020).

21. Potts, H. G., Stockdale, W. T. & Mommersteeg, M. T. M. Unlocking the secrets of the regenerating fish heart: Comparing regenerative models to shed light on successful regeneration. J Cardiovasc Dev Dis 8, 1–18 (2021).

22. Puhl, S. L. & Steffens, S. Neutrophils in Post-myocardial Infarction Inflammation: Damage vs. Resolution? Frontiers in Cardiovascular Medicine vol. 6 Preprint at 10.3389/fcvm.2019.00025 (2019).

23. Giles, A. J. et al. Dexamethasone-induced immunosuppression: Mechanisms and implications for immunotherapy. J Immunother Cancer 6, 1–13 (2018).

24. Bouvain, P. et al. Non-invasive mapping of systemic neutrophil dynamics upon cardiovascular injury. Nature Cardiovascular Research 2, 126–143 (2023).

25. Lai, S. L. et al. Reciprocal analyses in zebrafish and medaka reveal that harnessing the immune response promotes cardiac regeneration. Elife 6, 1–20 (2017).

26. Schubert, M. et al. Perturbation-response genes reveal signaling footprints in cancer gene expression. Nat Commun 9, (2018).

27. Besse, S., Nadaud, S., Balse, E. & Pavoine, C. Early Protective Role of Inflammation in Cardiac Remodeling and Heart Failure: Focus on TNFα and Resident Macrophages. Cells vol. 11 Preprint at 10.3390/cells11071249 (2022).

28. Fiordelisi, A., Iaccarino, G., Morisco, C., Coscioni, E. & Sorriento, D. NfkappaB is a key player in the crosstalk between inflammation and cardiovascular diseases. International Journal of Molecular Sciences vol. 20 Preprint at 10.3390/ijms20071599 (2019).

29. Zhang, Y. et al. Tumor Necrosis Factor-α and Lymphotoxin-α Mediate Myocardial Ischemic Injury via TNF Receptor 1, but Are Cardioprotective When Activating TNF Receptor 2. PLoS One 8, (2013).

30. Li, J. et al. B lymphocytes from early vertebrates have potent phagocytic and microbicidal abilities. Nat Immunol 7, 1116–1124 (2006).

31. Gao, J. et al. Novel functions of murine B1 cells: Active phagocytic and microbicidal abilities. Eur J Immunol 42, 982–992 (2012).

32. Martínez-Riaño, A. et al. Antigen phagocytosis by B cells is required for a potent humoral response. EMBO Rep 19, 1–15 (2018).

33. Jew, B. et al. Accurate estimation of cell composition in bulk expression through robust integration of single-cell information. Nat Commun 11, (2020).

34. Salmenov, R., Mummery, C. & Ter Huurne, M. Cell cycle visualization tools to study cardiomyocyte proliferation in real-time. Open biology vol. 14 240167 Preprint at 10.1098/rsob.240167 (2024).

35. Page, D. M. et al. An evolutionarily conserved program of B-cell development and activation in zebrafish. (2013) doi:10.1182/blood-2012-12.

36. Wienholds, E., Schulte-Merker, S., Walderich, B. & Plasterk, R. H. A. Target-Selected Inactivation of the Zebrafish rag1 Gene. Science (1979) (2002).

37. Petrie-Hanson, L., Hohn, C. & Hanson, L. Characterization of rag1 mutant zebrafish leukocytes. BMC Immunol 10, (2009).

38. Bevan, L. et al. Specific macrophage populations promote both cardiac scar deposition and subsequent resolution in adult zebrafish. Cardiovasc Res 116, 1357–1371 (2020).

39. Murray, P. J. et al. Macrophage Activation and Polarization: Nomenclature and Experimental Guidelines. Immunity vol. 41 14–20 Preprint at 10.1016/j.immuni.2014.06.008 (2014).

40. Vannella, K. M. & Wynn, T. A. Mechanisms of Organ Injury and Repair by Macrophages∗. Annual Review of Physiology vol. 79 593–617 Preprint at 10.1146/annurev-physiol-022516-034356 (2017).

41. Karra, R., Knecht, A. K., Kikuchi, K. & Poss, K. D. Myocardial NF-κB activation is essential for zebrafish heart regeneration. Proc Natl Acad Sci U S A 112, 13255–13260 (2015).

42. Bhattacharyya, S., Ghosh, S., Jhonson, P. L., Bhattacharya, S. K. & Majumdar, S. Immunomodulatory role of interleukin-10 in visceral leishmaniasis: Defective activation of protein kinase C-mediated signal transduction events. Infect Immun 69, 1499–1507 (2001).

43. Murray, P. J. & Wynn, T. A. Protective and pathogenic functions of macrophage subsets. Nature Reviews Immunology vol. 11 723–737 Preprint at 10.1038/nri3073 (2011).

44. Carey, C. M., Hollins, H. L., Schmid, A. V. & Gagnon, J. A. Distinct features of the regenerating heart uncovered through comparative single-cell profiling. Preprint at 10.1101/2023.07.04.547574 (2023).

45. Simkin, J. et al. Tissue-resident macrophages specifically express Lactotransferrin and Vegfc during ear pinna regeneration in spiny mice. Dev Cell (2024) doi:10.1016/j.devcel.2023.12.017.

46. Karin, M. & Clevers, H. Reparative inflammation takes charge of tissue regeneration. Nature 529, 307–3015 (2016).

47. Frangogiannis, N. G. The inflammatory response in myocardial injury, repair, and remodelling. Nature Reviews Cardiology vol. 11 255–265 Preprint at 10.1038/nrcardio.2014.28 (2014).

48. Horckmans, M. et al. Neutrophils orchestrate post-myocardial infarction healing by polarizing macrophages towards a reparative phenotype. Eur Heart J 38, 187–197 (2017).

49. Ma, Y. et al. Temporal neutrophil polarization following myocardial infarction. Cardiovasc Res 110, 51–61 (2016).

50. Soehnlein, O. & Lindbom, L. Phagocyte partnership during the onset and resolution of inflammation. Nature Reviews Immunology vol. 10 427–439 Preprint at 10.1038/nri2779 (2010).

51. Tokunaga, Y. et al. Comprehensive validation of T- and B-cell deficiency in rag1-null zebrafish: Implication for the robust innate defense mechanisms of teleosts. Sci Rep 7, (2017).

52. Jiao, J. et al. Regulatory B cells improve ventricular remodeling after myocardial infarction by modulating monocyte migration. Basic Res Cardiol 116, (2021).

53. Xu, Y. et al. Bone marrow-derived naïve B lymphocytes improve heart function after myocardial infarction: a novel cardioprotective mechanism for empagliflozin. Basic Res Cardiol 117, (2022).

54. Seifert, A. W. & Maden, M. New insights into vertebrate skin regeneration. in International Review of Cell and Molecular Biology vol. 310 129–169 (Elsevier Inc., 2014).

55. Deng, G. et al. Interleukin-10 promotes proliferation and migration, and inhibits tendon differentiation via the JAK/Stat3 pathway in tendon-derived stem cells in vitro. Mol Med Rep 18, 5044–5052 (2018).

56. Zhu, C., Yuan, T. & Krishnan, J. Targeting cardiomyocyte cell cycle regulation in heart failure. Basic Research in Cardiology vol. 119 349–369 Preprint at 10.1007/s00395-024-01049-x (2024).

57. Hinz, B. et al. The myofibroblast: One function, multiple origins. American Journal of Pathology 170, 1807–1816 (2007).

58. Meng, X. M., Nikolic-Paterson, D. J. & Lan, H. Y. TGF-β: The master regulator of fibrosis. Nature Reviews Nephrology vol. 12 325–338 Preprint at 10.1038/nrneph.2016.48 (2016).

59. Jeffery, W. R. Regressive evolution in astyanax cavefish. Annual Review of Genetics vol. 43 25–47 Preprint at 10.1146/annurev-genet-102108-134216 (2009).

60. Aspirasa, A. C., Rohnera, N., Martineaua, B., Borowskyb, R. L. & Tabina, C. J. Melanocortin 4 receptor mutations contribute to the adaptation of cavefish to nutrient-poor conditions. Proc Natl Acad Sci U S A 112, 9668–9673 (2015).

61. Lee, K. A., Wikelski, M., Robinson, W. D., Robinson, T. R. & Klasing, K. C. Constitutive immune defences correlate with life-history variables in tropical birds. Journal of Animal Ecology 77, 356–363 (2008).

62. Previtali, M. A. et al. Relationship between pace of life and immune responses in wild rodents. Oikos 121, 1483–1492 (2012).

63. O’Neill, L. A. J. & Pearce, E. J. Immunometabolism governs dendritic cell and macrophage function. Journal of Experimental Medicine vol. 213 15–23 Preprint at 10.1084/jem.20151570 (2016).

64. Buck, M. D., Sowell, R. T., Kaech, S. M. & Pearce, E. L. Metabolic Instruction of Immunity. Cell vol. 169 570–586 Preprint at 10.1016/j.cell.2017.04.004 (2017).

65. González-Rosa, J. M., Martín, V., Peralta, M., Torres, M. & Mercader, N. Extensive scar formation and regression during heart regeneration after cryoinjury in zebrafish. Development 138, 1663–1674 (2011).

66. Potts, H. G. et al. Discordant Genome Assemblies Drastically Alter the Interpretation of Single-Cell RNA Sequencing Data Which Can Be Mitigated by a Novel Integration Method. Cells 11, (2022).

67. Hao, Y. et al. Integrated analysis of multimodal single-cell data. Cell 184, 3573–3587.e29 (2021).

68. Hafemeister, C. & Satija, R. Normalization and variance stabilization of single-cell RNA-seq data using regularized negative binomial regression. Genome Biol 20, (2019).

69. Korsunsky, I. et al. Fast, sensitive and accurate integration of single-cell data with Harmony. Nat Methods 16, 1289–1296 (2019).

70. Zappia, L. & Oshlack, A. Clustering trees: a visualization for evaluating clusterings at multiple resolutions. Gigascience 7, (2018).

71. Farbehi, N. et al. Single-cell expression profiling reveals dynamic flux of cardiac stromal, vascular and immune cells in health and injury. (2019) doi:10.7554/eLife.43882.001.

72. Finak, G. et al. MAST: A flexible statistical framework for assessing transcriptional changes and characterizing heterogeneity in single-cell RNA sequencing data. Genome Biol 16, (2015).

73. Ashburner, M., et al. Gene Ontology: Tool for the Unification of Biology The Gene Ontology Consortium*. http://www.flybase.bio.indiana.edu (2000).

74. Korotkevich, G. et al. Fast gene set enrichment analysis. bioRxiv 060012 (2021) doi:10.1101/060012.

75. Love, M. I., Huber, W. & Anders, S. Moderated estimation of fold change and dispersion for RNA-seq data with DESeq2. Genome Biol 15, (2014).

76. Dobin, A. et al. STAR: Ultrafast universal RNA-seq aligner. Bioinformatics 29, 15–21 (2013).

77. Kolde, R. & Vilo, J. GOsummaries: An R Package for Visual Functional Annotation of Experimental Data. F1000Res 4, (2015).

78. 78. Warnes, G. R., et al. Package ‘gplots’ Title Various R Programming Tools for Plotting Data. (2016).

79. Stuart, T. et al. Comprehensive Integration of Single-Cell Data. Cell 177, 1888–1902.e21 (2019).

80. Korsunsky, I. et al. Fast, sensitive and accurate integration of single-cell data with Harmony. Nat Methods 16, 1289–1296 (2019).

81. Kassambara, A. ggpubr: ‘ggplot2’ Based Publication Ready Plots. (2023).

82. Liberzon, A. et al. Molecular signatures database (MSigDB) 3.0. Bioinformatics 27, 1739–1740 (2011).

83. Gentleman, R. C. et al. Open Access Bioconductor: Open Software Development for Computational Biology and Bioinformatics. Genome Biology vol. 5 http://genomebiology.com/2004/5/10/R80 (2004).

